# PKC-alpha promotes phosphorylation of KRAS suppressing its oncogenic properties

**DOI:** 10.1101/2022.05.24.493257

**Authors:** Tejashree Joglekar, Albert Ku, Ethan Schollaert, Yinan Gong, Jacob Stewart-Ornstein, Anatoly Urisman, Man-Tzu Wang

## Abstract

Oncogenic KRAS-driven cancers have long been considered as “undruggable” due to limited therapeutic options. While the recent success of KRAS-G12C inhibitors argues against the “undruggability” of KRAS, this treatment only benefits a small proportion of patients with KRAS mutant cancers, leaving an urgent need for modalities to target other KRAS mutants. KRAS-calmodulin (CaM) signaling axis reportedly regulates the oncogenic properties of KRAS through its C-terminal hypervariable region. Phosphorylation of KRAS by activated protein kinase C (PKC) uncouples KRAS-CaM, resulting in growth inhibition effective against the entire spectrum of KRAS hotspot mutations. However, broadly activating PKC could mediate tumor promoting signaling nodes and cause systemic toxicity, undermining its applicability as an anti-KRAS therapy. Here, we found that prostratin induces KRAS phosphorylation, resulting in an elevated level of active CaM in the cytosol of KRAS mutant cells, and consequentially suppresses their malignancies. A whole-genome wide CRISPR/Cas9 knockout screening, further confirmed by biochemical analysis, revealed that prostratin acts through activating PKCα. Functional studies confirmed PKCα as the sole kinase to phosphorylate KRAS and, therefore, a KRAS suppressor. Activation of PKCα induces senescence in KRAS mutant tumor cells through PTPN14, accompanied by a secretory phenotype contributing to the growth inhibition, and parallelly mediates a nuclear translocation of a CaM-dependent transcription activator, CAMTA-1, which can be a biomarker to indicate the activity of PKCα-KRAS-CaM axis. Our findings reveal a previously understudied regulation of KRAS-CaM axis by PKCα, which can be an actionable target for developing anti-KRAS therapeutics.

**One Sentence Summary:** This study deciphers a PKCα-led tumor suppressive effect specific to the “undruggable” KRAS-mutant tumor cells through the phosphorylation of KRAS and a consequently altered KRAS-CaM signaling axis.

## INTRODUCTION

Mutant RAS acts as a *bona fide* cancer driver in one-third of human cancers, generally accompanied by a dismal prognosis; consequently, mutant RAS is an attractive therapeutic target. The sequence of the first 165 amino acids in the three RAS isoforms, H-, N-, and K-RAS, whose mutations are directly linked to carcinogenesis, is highly homologous and determines their activity for signal transduction through canonical signaling pathways. However, the mutations in each RAS isoform show high tissue specificity suggesting that the different RAS isoforms may dictate distinct oncogenic outputs. KRAS is the most frequently mutated isoform among the three cancer-associated RAS family members and is preferentially linked to the three deadliest cancers, namely lung, colon, and pancreatic adenocarcinomas; thus, oncogenic KRAS is a major target in anticancer drug discovery [1, 2]. Because of its small size and compact structure, KRAS contains no accessible pocket where a drug can bind with high affinity, and thus no KRAS-specific inhibitors were approved for cancer treatment. Compounds targeting KRAS-G12C have recently entered clinical trials and showed promising response rates in patients with non-small cell lung cancers (NSCLCs) [3-7]. While this breakthrough validates KRAS as a strong target for intervention, some studies suggest that a subpopulation of KRAS mutant lung cancer cells can bypass the tumor-suppressive effects of KRAS-G12C inhibitors, leading to therapeutic resistance and potential recurrence [8]. Therapeutic approaches inhibiting other mutant forms of KRAS, such as G12D (the most common mutation in pancreatic ductal adenocarcinoma (PDAC) and in never-smoker patients with NSCLC), G12V, and Q61, remain unavailable [9-11]. Furthermore, a recent clinical trial has revealed that a subset of patients treated with KRAS-G12C inhibitor unfortunately acquired hotspot mutations in KRAS, including G12V and G12D [12]. New approaches are therefore urgently needed to suppress the malignancies driven by aberrancies across the mutational spectrum of KRAS.

Inhibition of canonical RAS effector downstream pathways, such as the MEK/ERK and PI3K/AKT pathways, has long been a popular alternative anti-KRAS approach and is undergoing clinical evaluation, but its efficacy and applicability has been limited by nonspecific cytotoxicity, incomplete inhibition of targeted effectors, and rebound activation of parallel signaling pathways [13, 14]. Therefore, identification of the regulatory activators or downstream signaling effectors specific and essential to the oncogenic output of KRAS can facilitate the design of inhibitors to suppress KRAS-driven malignancies with minimized unwanted systemic toxicity.

RAS proteins are expressed broadly across different anatomical regions despite the tissue-restricted spectrum of their mutations, e.g., KRAS mutations are prevalent in lung, colon, and pancreas cancers while NRAS mutations are predominantly found in melanoma. This suggests that mutations in particular RAS isoforms potentially dictate unique oncogenic features. Indeed, we previously defined that KRAS4B (abbreviated KRAS hereafter) promotes cancer stem cell (CSC)-like properties, an effect that cannot be reproduced by other RAS isoforms [15]. Importantly, these CSC-like properties cannot be reproduced in tumor models where an *HRAS* allele is knocked into the *KRAS* locus, suggesting that activating mutations in *KRAS* indeed translate into a distinct functional output in cancers and are not simply a reflection of a selective genetic predisposition influenced by gene-specific regulatory elements [15]. From the protein-structural point of view, only the flexible C-terminal hypervariable region (HVR) can distinguish KRAS from its closely related H- and N-RAS isoforms. Yet, little is known about whether the HVR domain dictates the oncogenic functions and signaling activity specifically mediated by KRAS. We further defined that the binding of KRAS to calmodulin (CaM) through its C-terminus determines the unique oncogenic properties of KRAS [15]. Phosphorylation of KRAS at serine 181, which is essential for this interaction, can dissociate the KRAS-CaM interaction and consequently block the oncogenic functions of KRAS in human cancers [15, 16].

Protein kinase C (PKC) has been shown to phosphorylate KRAS at serine 181 [17-19]. In the initial discovery, Bivona et al. have suggested that PKC agonist, specifically byrostatin, can suppress the growth of KRAS mutant cells and, in theory, be conveniently used as an anti-KRAS drug. Echoing their study, our previous study has used prostratin, another non-tumor-promoting PKC activator [19], to demonstrate that activating PKC can block the KRAS-CaM interaction and suppress oncogenic KRAS-driven malignancies [15]. However, the PKC family, which consists of 13 different isozymes, has long been considered oncoproteins due to the unambiguous tumor-promoting function of their potent ligands—phorbol esters—and the prevalence of PKC mutations in human cancers [20]. Indeed, activation of PKC promotes a broad range of signaling pathways involved in tumor promotion and/or acute systemic toxicity [21-23], diminishing the enthusiasm for developing PKC agonists as an anti-KRAS therapy.

Shifting the established paradigm, a recent study revealed unexpected loss-of-function (LOF) mutations in PKC and their tumor-suppressing functions in cancer cells [24]. In addition, LOF mutations in certain PKC isoforms show an association with oncogenic mutations in KRAS in human cancers [24], hinting a potential tumor-suppressor role of PKC in KRAS mutant cancers. In our prior study, while prostratin suppresses the malignancy of KRAS mutant pancreatic cancer cells both *in vitro* and *in vivo*, the pan-PKC activator phorbol 12-myristate 13-acetate (PMA) does not exhibit the same antitumor effects [15]. This observation suggests that prostratin, unlike PMA, may preferentially activate a PKC isozyme that specifically phosphorylates KRAS at serine 181. Identification and activation of the “KRAS-phosphorylating” PKC isozyme can therefore rewire the KRAS-mediated signaling pathway and consequently suppress its malignancies while minimizing the off-target effects on activation of tumor promoting signaling pathways and/or toxicity led by broad activation of PKC isozymes.

Here, we utilized prostratin that exhibited an unexpected anti-KRAS activity in our prior study to show that the phosphorylation of KRAS is specifically mediated by the activation of PKCα. Furthermore, pharmacological or genetical activation of PKCα induces a senescence program, including production of a specific secretome, via a CaM-dependent program. This study provides a rationale for targeting the PKCα-KRAS-CaM signaling axis to treat the malignancies driven by oncogenic KRAS.

## RESULTS

### Prostratin Induces KRAS Phosphorylation and Increases Cytoplasmic CaM Level

We previously showed that prostratin treatment can dissociate the intracellular KRAS-CaM interaction [15]. Disruption of the KRAS-CaM binding consequentially activated the downstream effector CaMKII, as indicated by its autophosphorylation level, and suppressed CSC-like properties uniquely mediated by oncogenic KRAS [15] (Fig. 1a). In this follow-up study, we first aim to validate that prostratin treatment directly induces the phosphorylation of KRAS at serine 181.

**Fig 1.**
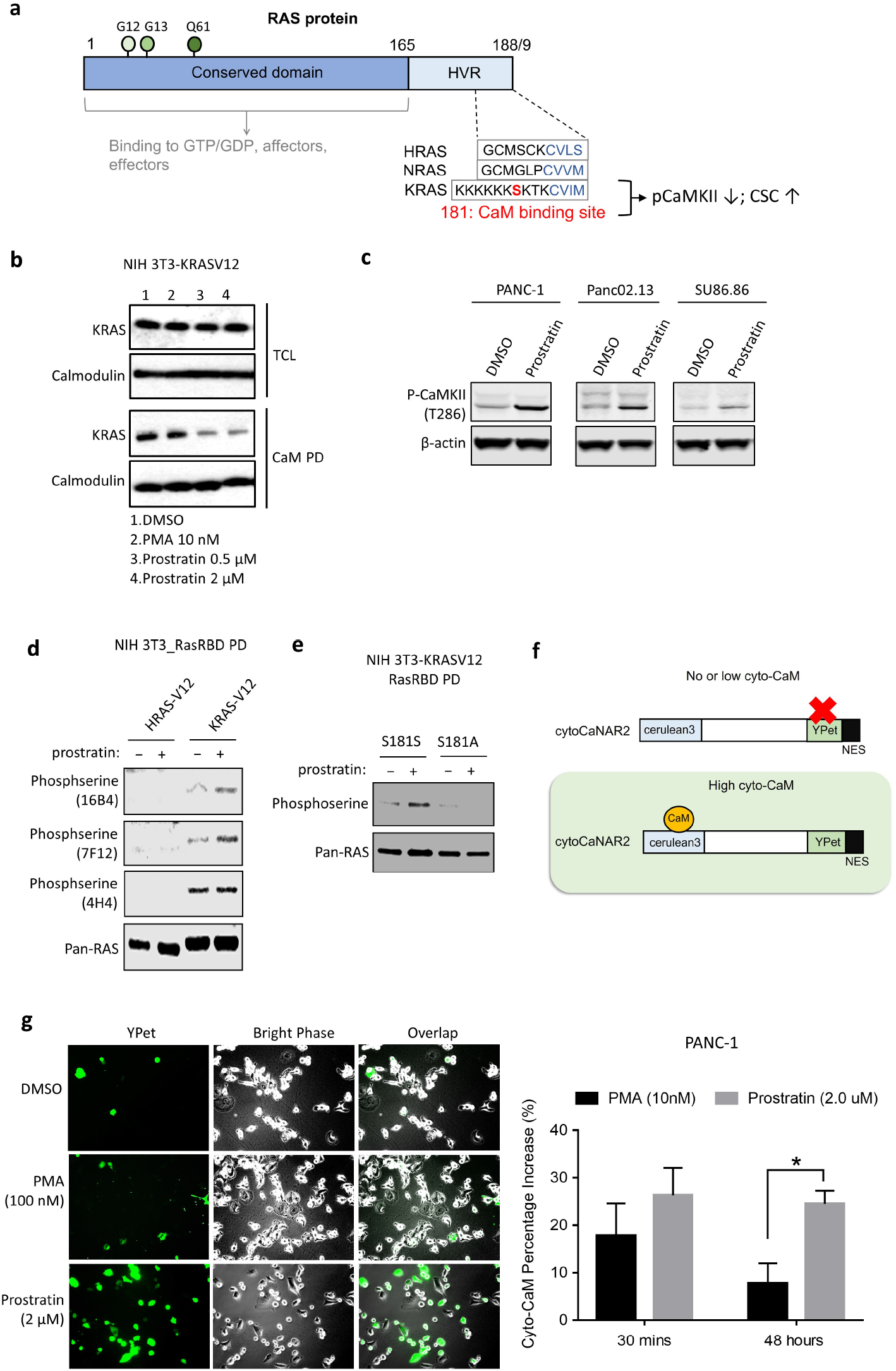
Prostratin induces phosphorylation of KRAS at serine 181, consequent KRAS-CaM dissociation, and increased phosphorylation of CaMKII. (a) Schematic of the RAS C-terminal HVR domain. The first 165 amino acids, which determine the binding of RAS proteins to GTP/GDP, regulatory factors, and signaling effectors, are highly homologous among RAS isoforms. Hotspot mutations of RAS proteins, including G12, G13, and Q61 alterations, occur in this region. KRAS-4B, herein abbreviated KRAS, has a unique HVR domain containing a polylysine sequence followed by a serine residue determining the binding to CaM. Phosphorylation of KRAS at serine 181 results in KRAS-CaM dissociation and consequently increases the phosphorylation of CaMKII and decreases cancer stem cell-like properties. (b) The CaM pulldown assay confirmed that prostratin, but not PMA, led to the KRAS-CaM dissociation in NIH 3T3 cells transformed with KRAS-G12V. TCL: total cell lysate. Cells were treated with each PKC activator for 48 hours before being lysed and subjected to the pulldown assay. (c) Western blot analysis of different KRAS mutant cancer cell lines treated with prostratin at 2 µM for 48 hours. Cells treated with prostratin but not those treated with bryostatin I or II showed increased phosphorylation of CaMKII. (d) Ras RBD pulldown followed by Western blotting for phosphoserine with different monoclonal antibodies. Prostratin treatment at 2 µM for 24 hours resulted in specific phosphorylation of KRAS at serine 181. The pulldown of active HRAS did not contain phosphorylated serine residues. (e) KRAS with the S181A mutation showed resistance to prostratin-induced phosphorylation. (f) Schematic illustrating the domain structures of the subcellularly targeted (cytoplasmic specific) variant of the CaNAR2 biosensor with a C-terminal nuclear export signal (NES). (g) Prostratin treatment increased the cytosolic distribution of active CaM in PANC-1 cells, as indicated by YPet activity from a cytosol-targeted FRET biosensor. The reading of YPet in each well before compound treatments was used as the baseline, and the YPet was measured again 30 minutes and 48 hours post-treatments. The percentage of increase in each well was calculated as (Reading YPet – baseline reading)/baseline reading * 100. The concentration of each compound is indicated in the figure. The representative pictures of increasing YPet positive cells in prostratin treated group are shown in the left. (n=4; * P < 0.05; ** P < 0.01) Means are plotted as bars with error bars showing s.d..

Since PKC activators exhibit a biphasic effect on the activation of PKC - i.e., they activate PKC to a lesser extent at higher concentrations [25], we first determined the working concentrations of prostratin for intracellular PKC activation. The well-studied non-tumor promoting bryostatin family, including bryostatin-I and -II, mediates activation of PKC by following a striking biphasic dose-response [26, 27] and therefore could be used as a control. ELISA that utilizes a specific synthetic peptide as a substrate for pan-PKC revealed that prostratin at 2 µM had the greatest abilities to activate PKC in mouse fibroblast and human cancer cell lines harboring different KRAS mutations (Supplementary Data Fig. 1a).

Reproducibly, prostratin but not PMA disrupted the binding of KRAS to CaM in transformed fibroblasts [15] (Fig. 1b) and increased the level of phosphorylated CaMKII in KRAS mutant cancer cells (Fig. 1b). Because no anti-phosphorylated KRAS (phospho-KRAS) antibody is available, and a poly-lysine sequence is located upstream of serine 181 in KRAS (Fig. 1a), which disables efficient trypsin digestion prior to mass spectrometric analysis, we used purified GST-Raf1 Ras-binding domain (RBD), glutathione agarose resin to pull down the active form of KRAS in cells and then performed Western blotting specific for phosphoserine. Prostratin treatment increased the level of phosphoserine in cells transformed with KRAS-G12V but not in those transformed with HRAS-G12V, which lacks a similar C-terminal domain for phosphorylation (Fig. 1d), or KRAS-G12V-S181A, which cannot be phosphorylated at serine 181 (Fig. 1e). These data suggest a unique ability of prostratin to induce phosphorylation of KRAS.

CaM is known to be abundantly present in various types of cells, leading to a potential caveat that the amount of CaM released from KRAS-bound form on the plasma membrane to cytosol may not be sufficient to modulate the subsequent CaM dependent signaling pathways. We then utilized a cytosolic calcineurin reporter [28], whose activity depends on the binding to CaM, to monitor the spatiotemporal subcellular patterns of active CaM upon the stimulation by prostratin (Fig. 1f). In this assay, the activity of a CaM dependent calcineurin reporter to indicate its activation in cytosolic fraction was significantly increased in the cells treated with prostratin when compared to other PKC activators (Fig. 1 g). These data suggest that the dissociation of KRAS-CaM by prostratin-induced KRAS phosphorylation can alter a downstream CaM-dependent signaling node.

Additionally, prostratin did not cause translocation of KRAS from the cell membrane to the cytoplasm or decrease the phosphorylation of ERK, further suggesting that its tumor-suppressive effect is not caused by loss of the cell membrane-associated form of KRAS (Supplementary Data Fig. 1b-c).

### Prostratin Selectively Inhibits Mutant KRAS-Driven Malignancies

In our previous report, we used prostratin as a proof-of-concept compound to demonstrate that the activation of PKC significantly prevents the initiation and inhibits the growth of pancreatic cancers, where mutant KRAS is the predominant driver [15]. In addition, such antitumor activity has been indicated to be specific to tumors initiated by oncogenic KRAS but not HRAS in the papilloma model [15].

However, it remained unknown whether prostratin also brings the growth-suppressing effect to other types of KRAS-mutant cancers, especially lung or colorectal adenocarcinomas where mutation in KRAS is one of the most common oncogenic events. To fill this knowledge gap, we tested multiple pancreatic, lung, and colon cancer cell lines, which harbor different mutant KRAS alleles, for their response to prostratin treatment to grow in 2D and 3D culture. The numbers of viable cells and single cell-derived colonies were significantly reduced upon short-term or long-term prostratin treatments (Fig. 2 a-b) in malignant cell lines that express different mutant forms of KRAS and display varied KRAS dependencies [29]. In addition, the suppressive effect of prostratin on colony formation was specific to KRAS-G12V-transformed cells and had no effect on the cells expressing KRAS-G12V-S181A, which cannot be phosphorylated at serine 181 (Fig. 2b, left panel), confirming our previous observation that the anti-KRAS activity of prostratin acts through the unique C-terminus of the KRAS protein. It has been increasingly recognized that under some circumstances, *in vitro* 2D and 3D culture systems can result in different bioactivities in cells [30, 31]. We therefore evaluated the effects of prostratin on the 3D growth of KRAS mutant cancer cells. Compared to the solvent control, prostratin significantly reduced the ability of KRAS-mutant cancer cells to form viable spheres in 3D culture (Fig. 2c, Supplementary Data Fig. 2a). Notably, human cancer cells expressing wild-type KRAS (Fig. 2c, left panel in dark grey circle). A mouse pancreatic adenocarcinoma cell line (iKRAS*), which maintains both wild-type KRAS alleles without doxycycline induction [32, 33], were insensitive to prostratin (Fig. 2b, right panel). iKRAS* cells with doxycycline induced expression of mutant KRAS acquired sensitivity to prostratin treatment (Fig 2b. right panel). Taken together, these results suggest that prostratin selectively suppresses the malignancy of cancer cells harboring constitutively activated KRAS independent of the mutational status in the G domain. Although the HVR domain, which may mediate sensitivity to prostratin, is identical in wild-type KRAS and mutant KRAS, the different responses of tumor cells harboring wild-type and mutant KRAS to prostratin treatment could result from the dependency on the oncogenic activity of KRAS. While most tumor cells expressing mutant KRAS rely heavily on its activity to maintain their malignancy, those in which wild-type KRAS is maintained can be dependent on other driver oncogenes for their survival and growth. Notably, prostratin exerted the greatest suppressive effect on the 2D and 3D growth of KRAS mutant cancer cells at 1 to 2 μM, the range of maximal function to activate pan-PKCs via detecting the phosphorylation of a synthetic peptide as a universal substrate (Supplementary Data Fig. 1a, Fig. 2a-c, Supplementary Data Fig. 2a).

**Fig 2.**
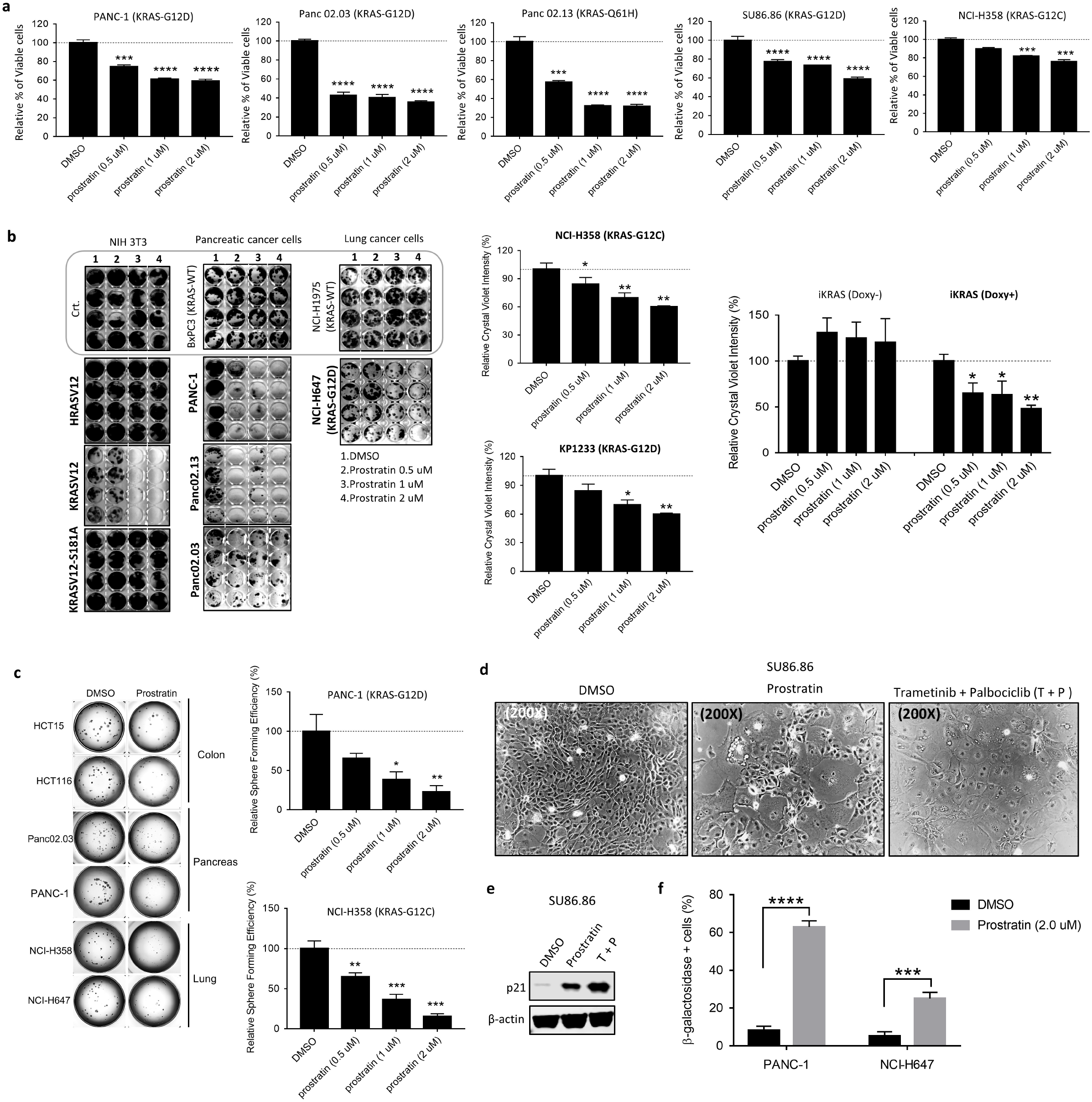
Prostratin suppresses the growth of KRAS mutant tumor cells via induction of cellular senescence but not apoptosis. (a) Upon treatments, prostratin reduced the numbers of viable human cancer cells when compared to DMSO treated control. The mutant forms of KRAS in each cell line are labeled as indicated. Cells were treated for 72 hours. A Cell Titer-Glo assay was used to determine cell viability, and the results were normalized to those in the control (DMSO-treated) group. (n=6; *** P < 0.001; **** P < 0.0001) Means are plotted as bars with error bars showing with standard deviation (s.d.). (b) Colony formation assay followed by crystal violet staining (left panel) in NIH 3T3 cells transformed with different RAS mutants, in human pancreatic cancer cell lines, and in human lung cancer cells in response to prostratin. The cell lines expressing wild type KRAS alleles are circled (left panel). The treatment concentrations are indicated. Cells were cultured in complete medium containing the compound until visible colonies formed. The crystal violet was dissolved, and the staining intensity was quantified by absorbance measurement (right panel). Prostratin blocked colony formation in NIH 3T3 cells transformed with KRAS-G12V and in cancer cell lines expressing mutant KRAS. NIH 3T3 cells expressing HRAS-G12V or KRAS-G12V-S181A maintained their colony-forming ability after prostratin treatment. The mutation status of each cell line is indicated, and the mutant forms of KRAS are labeled in bold font. Means of the absorbance at 570 nm with (n=4) are plotted as bars with error bars showing s.d.. (* P < 0.05, ** P < 0.01) (c) 3D sphere formation assay of a series of cancer cell lines harboring mutant KRAS and treated with prostratin. Prostratin treatment inhibited the sphere-forming ability of different human cancer cell lines. Representative images of a well containing spheres in a 96-well plate are shown in the left panel. In the representative images, cells were cultured in the completed media with Matrigel and 2 µM prostratin. DMSO was used as the solvent control. A 3D Cell Titer-Glo assay was used to quantitatively determine the overall viability of spheres. (n=6; * P < 0.05, ** P < 0.01; *** P < 0.001) Means are plotted as bars with error bars showing s.d.. (d) Representative images of SU86.86 cells exhibiting a senescent morphology after treatment with prostratin at 2 μM or with a combination of trametinib (10 nM) and palbociclib (0.1 μM) for 6 days. (e) Western blots indicating the expression level of p21 in SU86.86 cells treated with the indicated compounds for 6 days before the assay. (f) Prostratin treatment at 2 μM for 6 days significantly increased the percentage of β-Galactosidase positive cells quantified by flow cytometry in two different KRAS mutant cancer cell lines. (n=4; *** P < 0.001; **** P < 0.00001)

### Prostratin promotes senescence in KRAS-mutant cancer cells

Accumulating evidence suggests that activation of PKC may promote apoptosis [34, 35]. Indeed, Bivona et al. showed that activation of PKC by bryostatin slowed the growth of mouse tumors driven by oncogenic KRAS through induction of apoptosis [17]. However, no evidence indicated that prostratin treatment resulted in apoptosis in various prostratin-responsive cancer cells (Supplementary Data Fig. 2b). Instead, prostratin induced cellular senescence, defined by morphological change and senescence-associated β-galactosidase (SA-β-Gal) activity, in oncogenic KRAS-transformed human pancreatic epithelial cells, whereas immortalized cells or oncogenic HRAS-transformed cells showed no changes upon prostratin treatment (Supplementary Data Fig. 2c). Furthermore, KRAS-mutant cancer cells treated with prostratin exhibited a senescent morphology and phenocopied cells treated with both MAPK and CDK4/6 inhibitors (T+P), which can evidently reintroduce senescence in KRAS-mutant tumor cells [36] (Fig. 2d). Both prostratin treated- and T+P-treated cells showed increased p21 expression, which is one of the senescence-associated markers [37] (Fig. 2e). Flow cytometry-based SA-β-Gal activity assay further quantified and showed significantly increased senescent cell fraction in KRAS-mutant tumor cells treated with prostratin for at least 6 days (Fig. 2f). Prostratin-treated cells also showed increased p21 expression and reduced Lamin B1 expression, which is another senescence phenotype [38, 39] when compared to solvent- or byrostatin-treated cells (Supplementary Data Fig. 2d).

### PKCα is a Key Driver of Prostratin Sensitivity

The abovementioned results suggest that prostratin can be used to unbiasedly identify critical PKC isoforms or other potential targets involved in mediating KRAS phosphorylation. We performed a genome wide CRISPR/Cas9 knockout (KO) library screening. The human pancreatic cancer cell line PANC-1, which harbors KRAS-G12D and is sensitive to prostratin treatment, was used as the *in vitro* model for stable expression of the Cas9 protein and a lentiviral pooled CRISPR library containing 76,441 unique single-guide RNAs (sgRNAs). Cells expressing the CRISPR/Cas9 library were then treated with prostratin for 21 days. Knockout of a prostratin sensitivity driver gene could drive the resistance, survival, and proliferation of PANC-1 cells in the continuous presence of prostratin. In our genome wide screen, knockout of PKCα (encoded by *PRKCA*) strongly rescued survival, whereas other PRKCs were not identified as strong hits (Fig. 3a, labeled red). Notably, while *KRAS* and many other genes involved in canonical RAS pathways were also among the top 20 highest-ranking candidates (Fig. 3a, labeled blue), knockout of these genes, which are essential for the proliferation and survival of pancreatic tumor cells, may not directly contribute to prostratin resistance.

**Fig 3.**
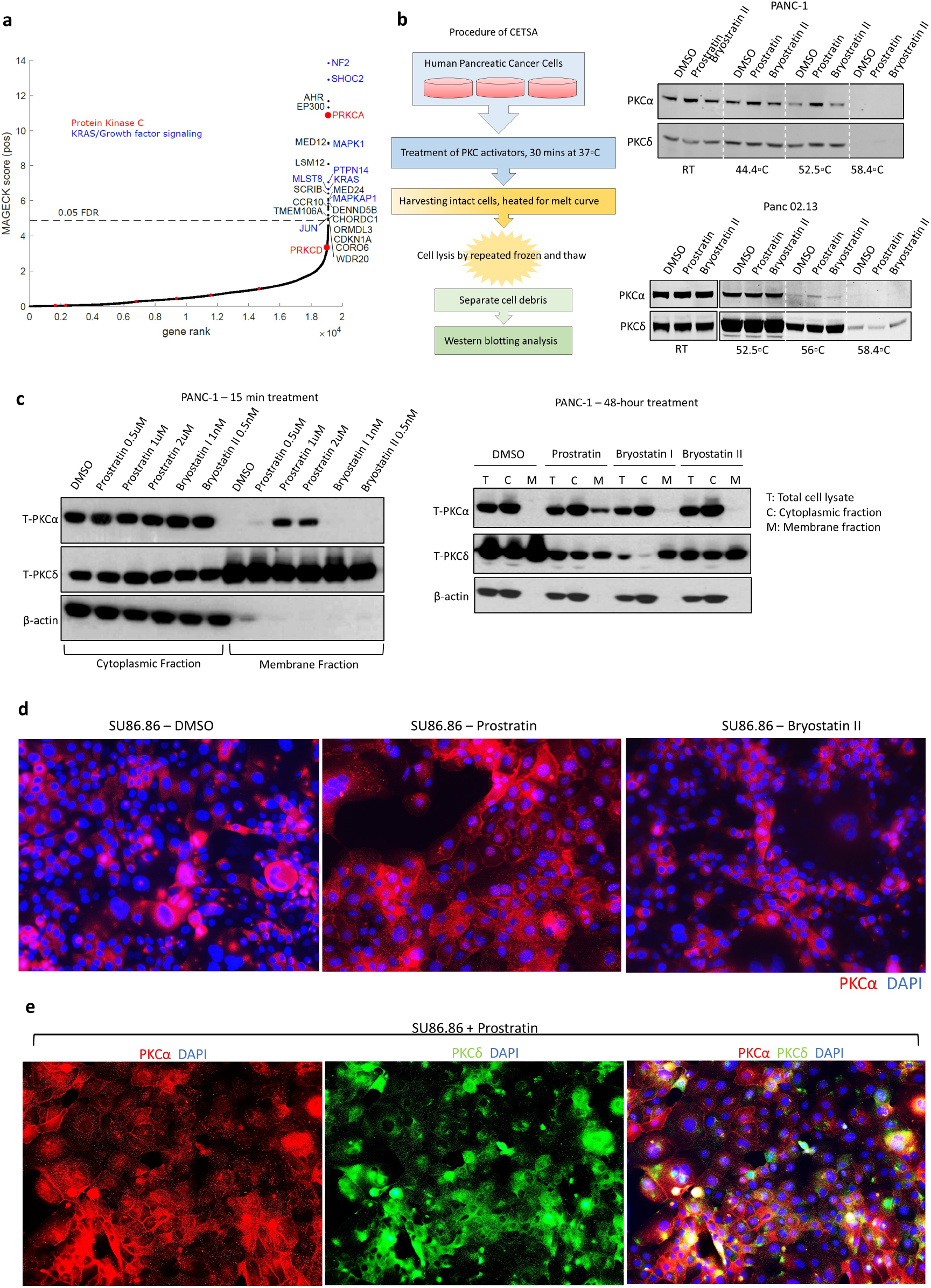
Prostratin binds and activates PKCα. (a) *In vitro* genome wide CRISPR/Cas9 knockout screen. PANC-1 were treated as described in the methods section. Plot showing the distribution of normalized CRISPR viability scores (prostratin treatment vs. DMSO control) for genes targeted by the sgRNAs in the library. The red dots indicate all members in the PKC family, and the blue dots indicate the genes involved in canonical RAS signaling pathways. *PRKCA*-targeting sgRNAs were negatively selected during prostratin treatment, suggesting that *PRKCA* is an essential gene to determine the survival of PANC-1 cells under prostratin treatment. The source data are provided as a Source Data file. (b) Scheme of the CETSA procedure used to examine the binding of PKC isozymes to ligands (prostratin or bryostatin) (left panel). CETSA followed by Western blot analysis suggested that prostratin had a better ability to bind and stabilize intracellular PKCα than did bryostatin II in two different KRAS mutant pancreatic cancer cell lines (right panel). PANC-1 cells were treated for 48 hours and Panc 02.13 were treated for 15 minutes. (c) Cell fractionation followed by Western blot analysis showed membrane translocation of PKCα in PANC-1 cells in response to prostratin (2 µM) but not bryostatin I (1 nM) or byrostatin II (0.5 nM) treatment. (d) Immunoflourescence staining for PKCα (indicated in red) in SU86.86 cells treated with prostratin at 2 µM or bryostatin II at 0.5 nM for 24 hours. Prostratin-treated cells exhibited visible membrane-localized PKCα, whereas those treated with bryostatin II exhibited cytoplasmic PKCα. (e) Immunoflourescence staining for PKCα (red) and PKCδ (green) in SU86.86 cells treated with prostratin at 2 µM for 24 hours.

### Prostratin Preferentially Binds and Activates PKCα

The PKC isozymes are classified into three groups: classical or conventional PKCs, including PKCα; novel PKCs, such as PKCδ; and atypical PKCs. All PKC members contain a conserved kinase domain, but only conventional and novel PKCs contain regulatory domains, which include C1 domains that act as ligand sensors and can bind to most PKC activators, including prostratin and bryostatin [40, 41]. We then investigated whether prostratin preferably binds to PKCα by using a cellular thermal shift assay (CETSA) [42], which allows us to investigate the thermal stability of proteins upon ligand binding in a cellular setting (Fig. 3b, left panel). Bryostatin is known to preferably stabilize and activate PKCδ [43, 44] and can therefore be used as a better control compound than PMA, which only activates PKC at the short term [45]. In the CETSA, prostratin had a better ability to bind PKCα than bryostatin II, whereas bryostatin II preferentially bound to and stabilized PKCδ in KRAS mutant cancer cells (Fig. 3b, right panel).

The hallmark of PKC activation is its translocation to cellular membranes in response to signal transduction through second messengers. We, therefore, examined whether the stabilization of PKCα by prostratin can directly translate into its activation via re-localization to the cellular membrane. The cell fractionation assay results suggested that prostratin specifically induced the translocation of PKCα from the cytoplasm to the cell membrane in two different KRAS mutant pancreatic cancer cells after short-term (15 minutes) and long-term treatment (48 hours) (Fig. 3c). On the other hand, bryostatin at long-term treatment promoted membrane translocation of PKCδ in the same cellular contents (Fig. 3c, right panel), consistent with the CETSA results in Fig. 3b. Immunofluorescence staining also confirmed that membrane translocation of PKCα was induced by prostratin but not by bryostatin II (Fig. 3d-e). Overall, these results suggest that prostratin and other PKC modulators bind and activate different PKC isoforms in KRAS mutant pancreatic cancer cells.

One potential caveat is that PKCα could be the most predominantly expressed PKC isozyme in KRAS mutant tumor cells and that its expression therefore accordingly affects the cellular responses to prostratin. *In silico* genome-wide transcriptome analysis in multiple tumor cell lines suggested that some tumor cells with PKCδ as the most dominant form of PKC remained highly sensitive to prostratin, further eliminating this possible caveat (Supplementary Data Fig. 3).

### KRAS Is a Substrate of PKCα

We next examined whether KRAS acts as a substrate of PKCα. Immunoprecipitation followed by Western blot analysis first confirmed that KRAS was one of the PKC substrates that can be phosphorylated by prostratin-activated PKC at serine residues (Fig. 4a). Furthermore, KRAS protein with a pseudo-phosphorylation mutation (aspartic acid (Asp)) or phosphorylation-resistant mutation (alanine (Ala)) in the CaM-binding domain (serine 181) no longer functioned as a PKC substrate (Fig. 4a). A direct interaction or binding is essential for kinases to activate their downstream substrates. To investigate the potential interactions between KRAS and PKC isozymes, we first utilized Rasless MEFs (mouse embryonic fibroblasts) with re-expression of either KRAS-G12V or NRAS-G12V. Ras RBD pulldown (PD) followed by Western blot analysis indicated that, in Rasless MEFs treated with prostratin, an interaction occurred between PKCα and active KRAS but not NRAS (Fig. 4b). Moreover, this interaction could not be reproduced in the cells treated with bryostatin II, and prostratin did not enhance the interaction between KRAS and PKCδ (Fig. 4b-c). Taken together, these results indicate that KRAS is a downstream substrate of PKCα. Co-transfecting HA-tagged-wild type or constitutively active PKCα [46] with oncogenic H- or -K-RAS in 293T cells, followed by immunoprecipitation of HA, further confirmed the specific interaction between PKCα and KRAS (Fig. 4d). These results suggest that the binding interaction of PKC with KRAS occurs in an isozyme-specific manner.

**Fig 4.**
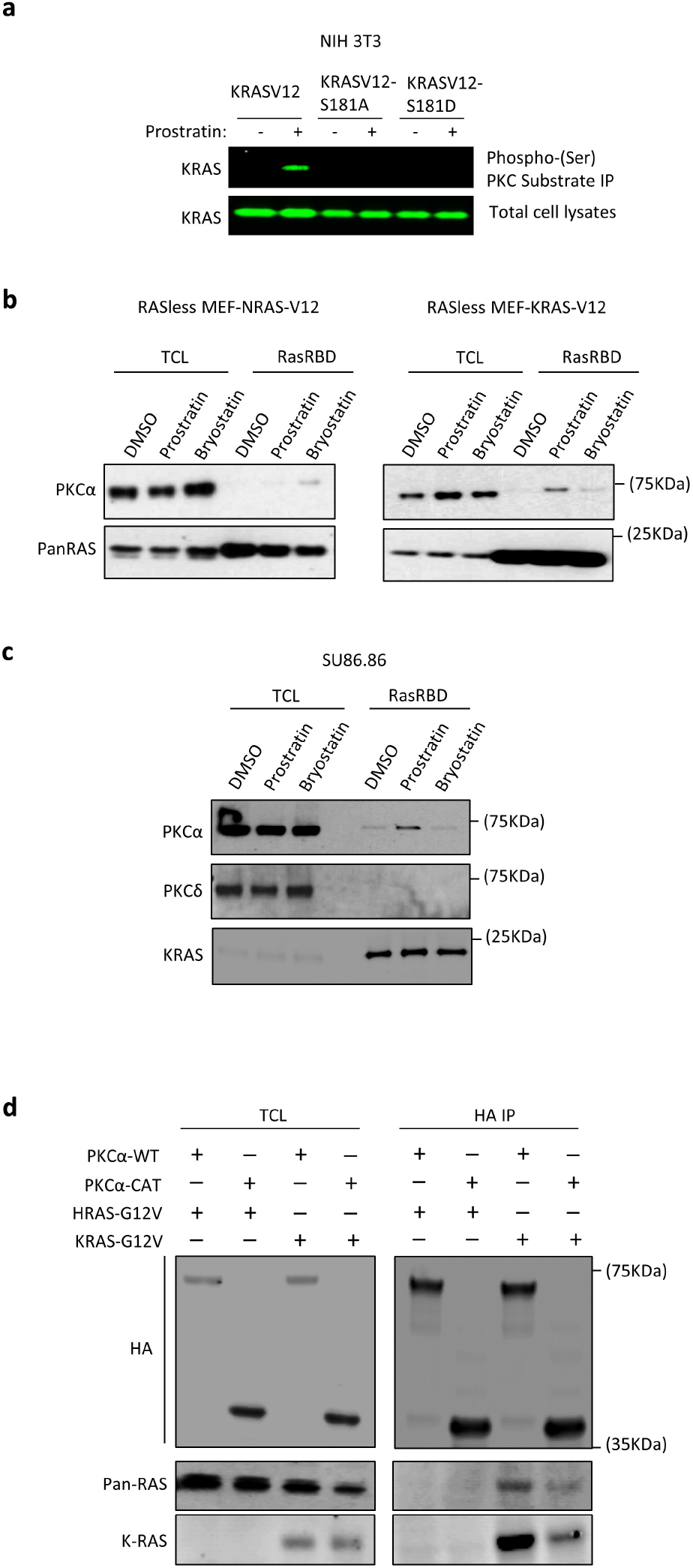
KRAS acts as a substrate of PKCα upon stimulation with prostratin. (a) Immunoprecipitation of phosphoserine PKC substrates revealed that KRAS was one of these substrates upon prostratin-induced activation. A point mutation at serine 181 of KRAS blocked this interaction. (b) RBD pulldown followed by estern blotting for phosphoserine indicated that KRAS but not NRAS interacts with and binds to PKCα in Rasless MEFs. Cells were treated with the indicated compounds for 24 hours before the pulldown assay (TCL: total cell lysate). (c) In an assay similar to that in (b), KRAS interacted with and bound to PKCα but not PKCδ in SU86.86 cells treated with prostratin at 2 µM for 24 hours. (d) Immunoprecipitation of HA tags in 293T cells co-transfected with HA-tagged PKCα expressing constructs and oncogenic RAS constructs at a 1:2 ratio suggested the interaction of PKCα and KRAS. Such interaction was absent between PKCα and HRAS. Due to the limitation of available antibody for probing HRAS post-immunoprecipitation, pan-RAS antibody that can detect all RAS isoforms was used to indicate the expression of HRAS.

### PKCα Is Essential for the Anti-KRAS Activity of Prostratin

To investigate the function of PKCα in mediating the ability of prostratin to suppress oncogenic KRAS, we established KRAS mutant pancreatic cancer cells in which PKCα was knocked out by CRISPR/Cas9, with or without restored expression (Fig. 5a). In this cell model, depletion of PKCα reduced the level of endogenous phospho-CaMKII compared to that in control cells expressing Cas9 alone, whereas restoring PKCα expression in the knockout clones increased the level of phospho-CaMKII (Fig. 5a). Furthermore, knocking out PKCα hindered KRAS-CaM dissociation caused by prostratin treatment in KRAS mutant cancer cells (Fig. 5b). In contrast, re-expression of PKCα restored KRAS-CaM dissociation in response to prostratin (Fig. 5b). Moreover, KRAS phosphorylation induced by prostratin in KRAS mutant cancer cells was diminished by a knockout of PKCα and was restored upon re-expression of PKCα (Fig. 5c). More importantly, loss of PKCα desensitized KRAS mutant tumor cells to prostratin treatment in terms of survival and colony formation, and sensitivity to prostratin was restored in cells with restored expression of PKCα (Fig. 5d, Supplementary Data Fig. 4a).

**Fig 5.**
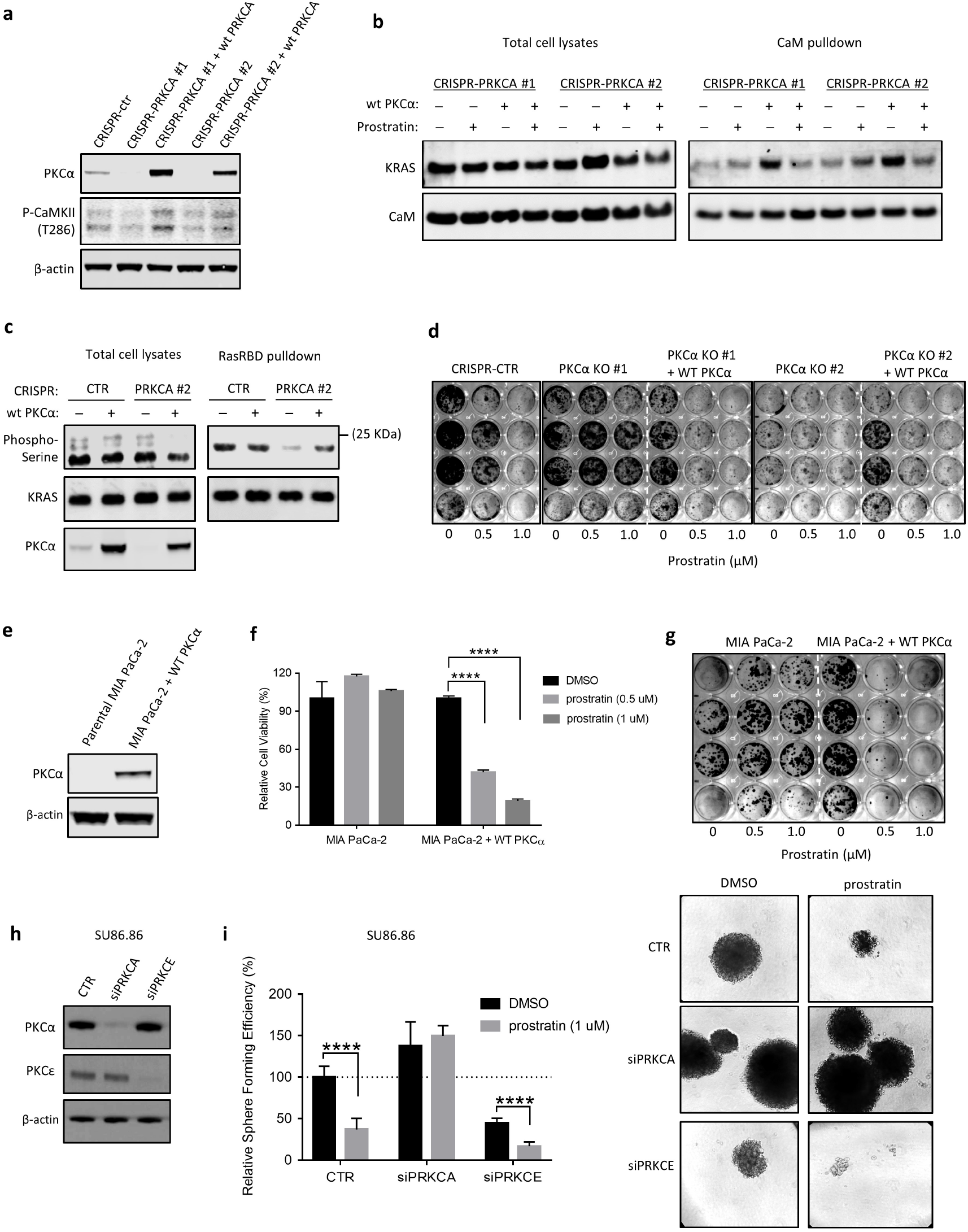
The expression of PKCα determines the responsiveness of KRAS mutant cancer cells to prostratin. (a) Western blot analysis confirmed knockout of PKCα by CRISPR/Cas9 and re-expression of PKCα by re-introduction of wild-type PKCα (Horizon clone #OHS5899-202625016) in PANC-1 cells. (b) CaM pulldown followed by Western blotting suggested that the KRAS-CaM interaction was not disrupted upon prostratin treatment in PANC-1 cells with PKCα knockout. Cells with re-expression of PKCα responded to prostratin and showed KRAS-CaM dissociation. Cells were treated with prostratin at 2 µM for 24 hours before the pulldown assay. (c) Ras RBD pulldown followed by Western blotting for phosphoserine suggested loss of endogenous phosphorylated KRAS in PANC-1 cells with PKCα knockout. Re-expression of PKCα restored the phosphorylation of KRAS in PANC-1 cells. (d) A colony formation assay with crystal violet staining was used to evaluate the effects of prostratin treatment on the long-term survival and proliferation of PANC-1 cells with PKCα knockout and with or without re-expression of PKCα. Cells were seeded at 100 cells per well and cultured in complete medium with the indicated compounds for 14 days until visible colonies formed. (e) Western blot analysis confirmed the overexpression of PKCα in MIA PaCa-2 cells, which do not have endogenous expression of PKCα. (f) CellTiter-Glo assay to evaluate the viability of MIA PaCa-2 cells with or without PKCα overexpression after treatment with prostratin for 72 hours. (n=6; **** P < 0.0001) Means are plotted as bars with error bars showing s.d.. (g) Colony formation assay followed by crystal violet staining in MIA PaCa-2 cells treated with prostratin. (h) Western blot analysis confirmed the siRNA-knockdown of PKCα (pool, Horizon #L-003523-00-0005) or PKCε (pool, Horizon #L-004653-00-0005) in SU86.86 cells. (i) 3D sphere formation assay in SU86.86 cells with knockdown of PKCα or PKCε. (Right panel) The representative images of spheres (image power = 200X). (Left panel) The viable spheres were quantified by 3D CellTiter Glo assay and scramble siRNA transfected cells with DMSO treatment were used as control for normalization. (n=6; **** P < 0.0001) Means are plotted as bars with error bars showing s.d..

Among all the cancer cell lines harboring mutant KRAS, MIA PaCa-2 and Panc03.27, with no detectable endogenous PKCα expression, were outliers and did not respond to prostratin treatment (Supplementary Data Fig. 4b-c). Overexpression of PKCα significantly sensitized MIA PaCa-2 cells to prostratin (Fig. 5e-g). Next, we aimed to eliminate the possibility that PKCδ, as the most dominantly expressed form of PKC in KRAS mutant cancer cells [47](Supplementary Data Fig. 3b), also phosphorylates KRAS. Since PKCδ is required for survival of pancreatic and lung cancer cells [48-50], knocking out its expression by CRISPR/Cas9 induced massive cell death of the tumor cells (data not shown). We therefore used different shRNA hairpins to knock out PKC-α or -δ (Supplementary Data Fig. 4d). Under comparable knocking down efficiency (∼60%), the cells with knockdown of PKCδ retained the response to prostratin and showed reduced interaction of KRAS-CaM (Supplementary Data Fig. 4 d-e), increased phosphorylation of KRAS (Supplementary Data Fig. 4f) and reduced long-term growth (Supplementary Data Fig. 4g) upon prostratin treatment. Furthermore, in SU86.86 cells, which express detectable PKCα and PKCε (Supplementary Data Fig. 3b), knocking down PKCε by siRNA did not alter the cellular response to prostratin treatment, whereas the cells with knockdown of PKCα were resistant to the treatment of prostratin in 3D culture (Fig. 5h-i). Taken together, these results confirmed that the intact expression of PKCα, but not other cancer-associated PKC isozymes, is functionally important for prostratin-induced KRAS phosphorylation and the consequent suppression of malignancy in KRAS mutant cancer cells.

### PKCα suppresses KRAS-mediated malignancies

Introduction of an HA-tagged constitutively active mutant of PKCα (PKCα-CAT) [46] into PANC-1 cells with PKCα knockout by CRISPR/Cas9 increased the phosphorylation of CaMKII and suppressed colony formation even in the absence of prostratin (Supplementary Data Fig. 5a; Fig. 6a). The growth inhibition caused by the ectopic expression of PKCα-CAT only occurred in the KRAS mutant tumor cells, but not in the tumor cells harboring wild type KRAS (BxPC3 cells) or mutant NRAS (NCI-H1299 cells) (Supplementary Data Fig. 5 b-c), resembling the selective suppressive effect of prostratin in tumor cells expressing mutant KRAS. Re-expression of wild type PKCα in MIA PaCa2 cells, which exhibit no detectable endogenous PKCα, significantly suppressed their growth as xenograft tumors (Supplementary Data Fig. 5d) and knocking out PKCα by CRISPR/Cas9 dramatically increased the growth of PANC-1 xenograft tumors (Fig. 6b), indicating the tumor suppressive role of PKCα in KRAS mutant tumor cells. In addition, knocking out PKCα increased the tumor initiation frequency in PANC-1 cells when transplanted at lower numbers of tumor cells (0.1 × 10^6^ per flank) than the routinely used numbers (reported 1 to 10 × 10^6^) (Fig. 6b), indicating that the expression of PKCα may suppress CSC-like properties in KRAS mutant tumor cells. Overexpression of wild type PKCα or constitutively active PKCα significantly increased cytosol-localized active CaM (Fig. 6c). These results suggest that activated PKCα specifically suppresses the malignancies of KRAS mutant tumor cells and confirm that active PKCα recapitulate the tumor suppressive effect of prostratin.

**Fig 6.**
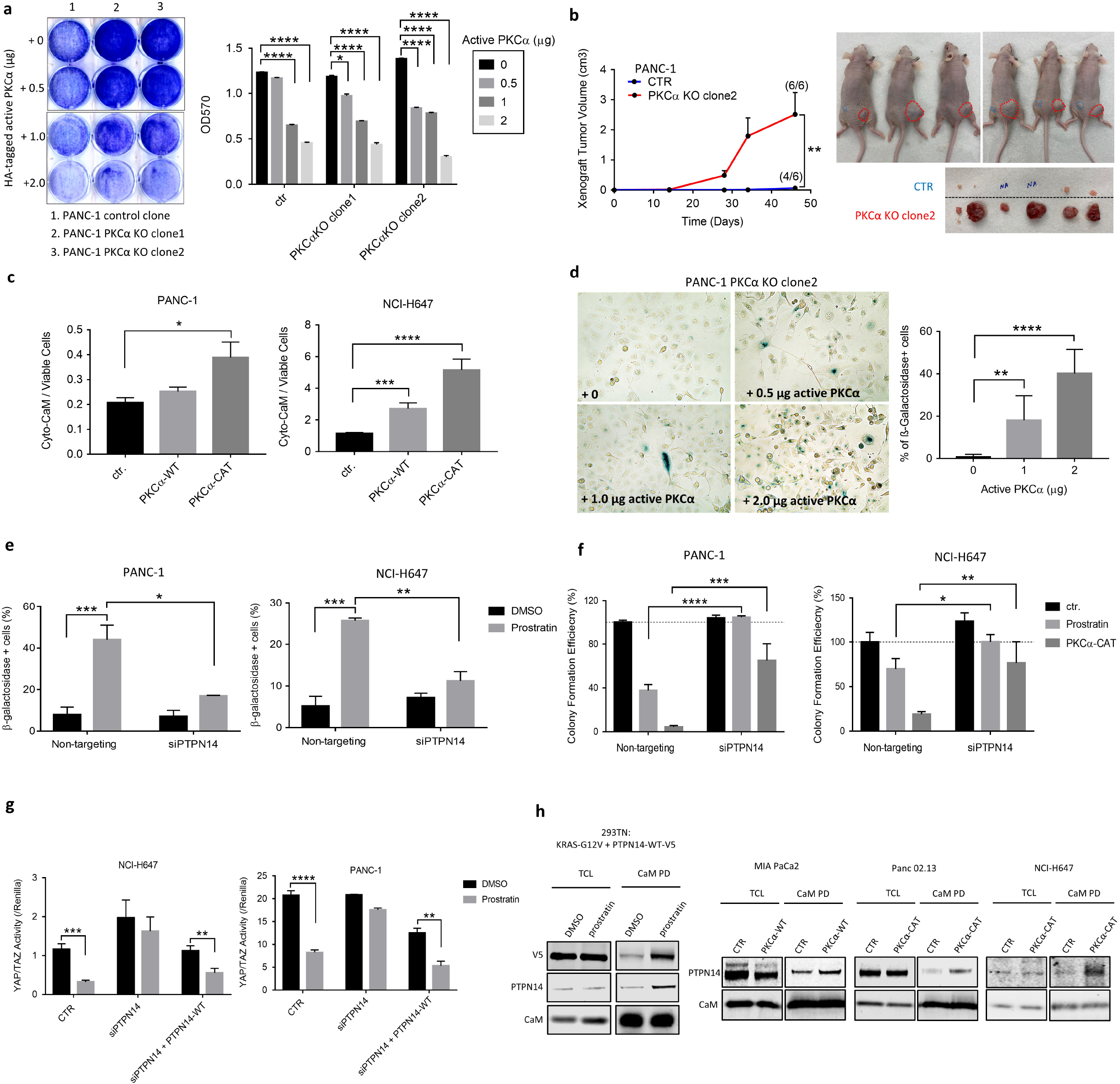
Over-expression of PKCα or expressing constitutively active PKCα inhibits the growth and induces senescence in KRAS mutant cancer cells. (a) Colony formation assay with visualization by crystal violet staining showed the inhibitory effect of constitutively active PKCα on the long-term growth of PANC-1 cells. The same number of cells was seeded in 6-well plates, and the cells were then transfected with constitutively active PKCα. Twelve hours post transfection, the cells were returned to complete medium for culture. The cells were stained for crystal violet on the 10^th^ day post transfection. The experiment was repeated for four times, the crystal violet was dissolved, and the intensity of staining was quantified at absorbance 570 nm. (n=4; **** P < 0.0001). Means are plotted as bars with error bars showing s.d.. (b) Tumor growth curve of xenograft tumors derived from PANC1 cells. Error bars here indicate standard error of the mean (SEM). (n=6; ** P < 0.01) The images of mice bearing subcutaneous tumors are shown on the right. Tumors derived from control cells were circled in blue, and tumors from knockout group were circled in red. The images of actual and visible tumors are shown on the bottom right. (c) Cytosolic distribution of active CaM, indicated as YPet activity from a cytosol-targeted FRET biosensor, in the cells co-transfected with biosensor and PKCα expressing constructs. The YPet in each well was measured 48 hours post-transfection, and the reading was normalized by the number of viable cells per well. (n=4; ** P < 0.01; *** P < 0.001; **** P < 0.0001) Means are plotted as bars with error bars showing s.d.. (d) β-Galactosidase staining was used to identify PANC-1 cells undergoing senescence expressing different levels of constitutively active PKCα. The cells were stained on the 6^th^ day post transfection of active PKCα. The number of senescent cells was counted manually in a double-blind manner. (n=4; ** P < 0.01; **** P < 0.0001) (e) Percentage of β-galactosidase positive cells, stained with CellEvent™ Senescence Green and measured by flow cytometry analysis, in the KRAS mutant tumor cells. The cells were first transfected with siRNA against *PTPN14* and treated with 2 μM prostratin for 5 days to induce senescence. Non-targeting siRNA was used as control. (n=3; * P < 0.05; ** P < 0.01; *** P < 0.001) (f) Colony formation assay followed by crystal violet staining suggested that knocking down PTPN14 by siRNA rescued the growth inhibition caused by activation of PKCα in KRAS mutant cancer cells. The cells were transfected with siRNAs, followed by treatment of prostratin at 2 μM or transfected with constitutively active PKCα. The cells were allowed to grow for 10 days or until visible colonies formed. The cells were then fixed and stained with crystal violet, later dissolved in 1% SDS for further determination of colony density. (n=4; * P < 0.05; ** P < 0.01; *** P < 0.001; **** P < 0.0001) Means are plotted as bars with error bars showing s.d.. (g) YAP/TAZ transcriptional activity indicated the firefly luciferase suggested that prostratin treatment significantly reduced the YAP activity in two different KRAS mutant cancer cell lines. Knocking down PTPN14 diminished such effect, which could be rescued by restored expression of PTPN14. The cells were first transfected with siPTPN14. 18 hour-post transfections of siRNA, the cells were co-transfected with YAP/TAZ reporter and Renilla luciferase construct at 2:1 ratio. 24 hours post transfection, the cells were treated with prostratin at 2 µM for 24 hours before being harvested for dual luciferase assay. (n=4; ** P < 0.01; **** P < 0.0001) (h) CaM pulldown in different cell lines to indicate the interaction between CaM and PTPN14, which was enhanced by the treatment of prostatin at 2µM for 24 hours. (TCL: total cell lysate; PD: pulldown)

### Active PKCα is downregulated in progressed murine pancreatic tumors

The results above suggest activation of PKCα as a tumor suppression mechanism in KRAS mutant tumor cells, leading to a question whether deactivation or lose of PKCα is a key step to facilitate tumorigenesis driven by oncogenic KRAS. Since loss of *PRKCA* gene is not known to be associated with mutant KRAS driven tumorigenesis, we investigated whether a potential deactivation of PKCα occurs with progression of KRAS-driven cancers. To evaluate the activation of PKCα during oncogenic KRAS-induced tumorigenesis, we turned to a well-studied pdx-Cre; KRAS-G12D pancreatic cancer mouse model. To increase the tumor-occurrence rate and facilitate tumor progression in our model, we administered caerulein at 50 μg/kg, via eight intraperitoneal (I.P.) injections over 48 hours, to induce pancreatitis in pdx-Cre; KRAS-G12D mice ≤ 8 weeks old. Consistent with the reported literature, 100% of our pdx-Cre; KRAS-G12D mice without loss of p53, when subjected to caerulein induced pancreatitis, rapidly developed distinct grades of PanINs [51, 52] (Supplementary Data Fig. 5e). Phosphorylation of PKCs is important for their function and/or used as an indication of their activation. The level of phosphorylated PKCα at serine 657 within the hydrophobic motif which is necessary for the stability of active PKC protein [53], was reduced in the high-grade pancreatic tumors (PanIN-3) developed in the mice at 7 months old when compared to the low-grade pancreatic tumors (PanIN-1) (Supplementary Data Fig. 5f), indicating the potential downregulation and inactivation of PKCα during the development and progression of oncogenic KRAS-driven pancreatic cancer.

### Activated PKCα induces senescence in KRAS mutant tumor cells

Resembling the effects of prostratin treatment, ectopic expression of constitutively active PKCα suppressed the growth of KRAS mutant cancer cells by inducing cellular senescence, as indicated by SA-β-Gal staining (Fig. 6d). Among all the key mediators of cellular senescence, such as p16, TP53, and p21, the expression of p21 at the protein level was increased in a series of KRAS mutant tumor cells, either treated with prostratin or expressing active PKCα (Supplementary Data Fig. 6a-b). Such changes in p21 expression also occurred at the transcriptional level (Supplementary Data Fig. 6c). Cell fractionation assay revealed that there was more nucleus-localized p21, as an indication of functional p21, in the tumor cells expressing active PKCα than control cells (Supplementary Data Fig. 6d). These data suggest that activation of PKCα suppresses the growth of KRAS mutant tumor cells through induction of cellular senescence. Furthermore, the treatment with a CaM inhibitor, W7, blocked the increase in p21 expression caused by the pharmacologically activated PKCα in human pancreatic cancer cells (Supplementary Data Fig. 6e), suggesting that CaM is involved in the cellular senescence triggered by activated PKCα in the KRAS mutant tumor cells. In addition, KRAS mutant tumor cells with knockdown of PKCδ retained the response to prostratin and underwent cellular senescence after long term of treatment (>5 days), whereas the treatment of prostratin did not induce senescence in the tumor cells with knockdown of PKCα (Supplementary Data Fig. 6f).

### PTPN14 mediates the PKCα induced senescence via suppression of YAP

Identifying the key downstream effector to mediate PKCα-induced senescence in KRAS mutant cells may enable us to design alternative anti-KRAS therapeutic strategies to recapitulate the effects of activating PKCα while minimizing the potential tumor promoting effects driven by PKCα. Thus, we used siRNAs to knock down top 5 genes, excluding PKCα, KRAS, and other canonical RAS signaling nodes, from our genome-wide KO library screening (Fig. 3a). We also included p38 MAPK (*MAPK11*) because activation of p38 MAPK signaling has been reported to be involved in PKCα-mediated tumor suppression and cellular senescence in oncogenic KRAS-induced lung cancer [54]. With comparable knockdown efficiency (Supplementary Data Table 1), only depletion of PTPN14 compromised the senescence program including increased senescence associated β-galactosidase positive population (Fig. 6e; Supplementary Data Table 1) and p21 expression (Supplementary Data Fig. 6g) in KRAS mutant cancer cells with intact endogenous PKCα (PANC-1) or with restored expression of PKCα (MIA PaCa-2). Furthermore, knocking down PTPN14 by siRNA rescued the inhibited colony formation ability of KRAS mutant cancer cells caused by activated PKCα (Fig. 6f). These data suggest that PTPN14 is a downstream mediator of PKCα-induced tumor suppression in KRAS mutant cancers.

Digging deeper into the mechanism of how PKCα mediates its tumor suppressive effects, our results above indicate that genetic or pharmacological activation of PKCα can induce a senescence program through PTPN14. PTPN14 reportedly interacts with the pivotal effector of Hippo signaling pathway, YAP, and further inhibits its transcriptional function [55, 56]. In addition, PTPN14-YAP signaling axis has been reported to be integral to p53-mediated tumor suppression in pancreatic cancer [56, 57], potentially coordinating multiple cell fates, including senescence [58]. YAP/TAZ-responsive luciferase reporter assay revealed that prostratin treatment significantly reduced the activity of YAP in KRAS mutant cancer cells yet was ineffectual to the cells where PTPN14 had been knocked down by siRNA (Fig. 6g). Re-expressing wild type PTPN14 in the cells with knockdown of PTPN14 further restored their response to prostratin and showed reduced the transcriptional activity of YAP (Fig. 6g).

Even though the mechanism underlying the regulation of PTPN14 by KRAS-CaM axis remains to be further clarified, Calmodulation database and meta-analysis predictor [59] predicts that PTPN14 has multiple CaM interacting motifs (Supplementary Data Table 2) and likely to be a CaM binding protein, raising the possibility that the binding to CaM determines the function of PTPN14. CaM pulldown (CaM PD) further suggested a direct physiological interaction between CaM and PTPN14 in 293T cells which expressed KRAS-G12V and V5-tagged wild type PTPN14 (Fig. 6h, left panel). In addition, the overexpression of PKCα or ectopic expression of constitutively active PKCα further enhanced the binding of PTPN14 to CaM in human pancreatic cancer cells with mutant KRAS (Fig. 6h, right panel).

### CAMTA1 is a bio-indicator of PKCα/KRAS/CaM axis

Based on the data above, phosphorylation of KRAS by PKCα releases active CaM from the KRAS-CaM complex into the cytosol, sequentially modulating a broad range of cellular functional outputs, including cellular senescence. It has been previously reported that CaM and certain CaM-dependent signaling pathways positively regulate senescence programs in plant cells [60-62]. In particular, the CaM binding transcription activators (CAMTAs) are a group of transcription factors that act as target proteins of CaM and are widely conserved across eukaryotes [63]. In plants, the CAMTAs function significantly in growth and development, defenses against biotic stress, and senescence in the response to abiotic stresses [64]. Here, our nucleus extraction followed by Western blotting analysis revealed that, between the two CAMTA proteins encoded in mammals, CAMTA-1 was less localized in the nuclear fraction from the cells transformed by KRAS-G12V when compared to the ones expressing HRAS-G12V or KRAS-G12V-S181D mutant, which are incapable of CaM binding (Supplementary Data Fig. 6h, left panel). Overexpression of PKCα increased both CAMTA proteins in the nuclear fraction of MIA PaCa-2 cells (Supplementary Data Fig. 6h, middle panel). Prostratin treatment reinforced the expression of CAMTA-1 in the nuclei of PANC-1 cells, whereas the combination of Trametinib and Palbociclib that induces cellular senescence through co-inhibition of CDK4/6 and MEK did not bring the same effect (Supplementary Data Fig. 6h, right panel). This result indicates that the increase of nucleus-localized CAMTA-1 is specific to the cells with activated PKCα. Notably, in the Lrig1-Cre/ER/LSL/KRAS-G12D mice where tamoxifen-inducible Cre recombinase initiates the expression of KRAS-G12D and consequential formation of papilloma during wound-healing [15], administrating therapeutic level of prostratin not only prevented the tumor initiation [15] but also significantly elevated the expression level of nucleus-localized CAMTA-1 if the tumors formed (Supplementary Data Fig. 6i), thus validating that CAMTA-1, as a CaM responder, can be potentially used as a biomarker to indicate the anti-KRAS therapeutic efficacy of PKCα agonists *in vivo*.

### Activation of PKCα induces a secretory phenotype in KRAS mutant tumor cells

Senescent cells are trapped in a prolonged growth arrest yet remain metabolically viable and biologically active through secretion of various cytokines and chemokines, also known as senescence-associated secretory phenotypes (SASPs) [65]. SASPs reportedly have physiological and pathological roles in aging-related diseases and cancers [66]. We then evaluated the secretory proteins generated by the KRAS mutant cells undergoing active PKCα-induced senescence. Analysis of an antibody array, which detects 36 different human cytokines or chemokines, revealed that the expression of constitutively active PKCα elevated the expression of multiple secretory proteins in KRAS mutant pancreatic cancer cells (Fig. 7a). Among the analyzed cytokines or chemokines, the expressions of CCL3/4 (MIP-1α/-1β) and CXCL-1 were consistently changed (fold change in intensity was greater than 3) across two different cell lines upon genetic activation of PKCα (Fig. 7a). The expressions of the same chemokines were also increased in the cells treated with prostratin when compared to the senescence-inducing treatment with Trametinib and Palbociclib (Extended Data Fig. 7a), indicating that CCL3/4 (MIP-1α/-1β) and CXCL-1 might constitute a unique SASP caused by the activation of PKCα in KRAS mutant tumor cells. ELISA confirmed that, upon genetic or pharmacological activation of PKCα, different lung and pancreatic cancer cells harboring mutant KRAS produced and secreted CXCL1 when compared to the control cells (Fig 7b; Supplementary Data Fig. 7b). Notably, the increased expression of *CXCL1* caused by activated PKCα occurred early at the transcriptional level, and inhibition of CaM activity by W7 treatment or knockdown of PTPN14 suppressed the increase in *CXCL1* expression in the KRAS mutant tumor cells upon PKCα activation (Supplementary Data Fig. 7c-d). In addition, blocking the activity of CXCL1 by neutralizing antibody (Fig. 7b) partially rescued the growth inhibition caused by activated PKCα in KRAS mutant tumor cells (Fig. 7c; Supplementary Fig. 7g). Despite being non-detectable or low as the secreted forms in some cells (i.e. PANC-1 and NCI-H647), CCL3 and CCL4 were highly secreted from Panc02.13 cells with activation of PKCα (Supplementary Data Fig. 7e-f). In Panc02.13 cells, neutralizing the activity of CCL3 or CCL4 by antibodies (Fig. 7c, right panel) resulted in a similar rescued colony formation as neutralization of CXCL1 (Fig. 7c, left panel). These results suggest that the production of MIP-1α, -1β and/or CXCL-1 are a SASP specific to the KRAS mutant tumor cells undergoing senescence induced by altered PKCα-KRAS-CaM signaling axis, which contributes to the growth inhibition caused by PKCα-induced senescence.

**Fig 7.**
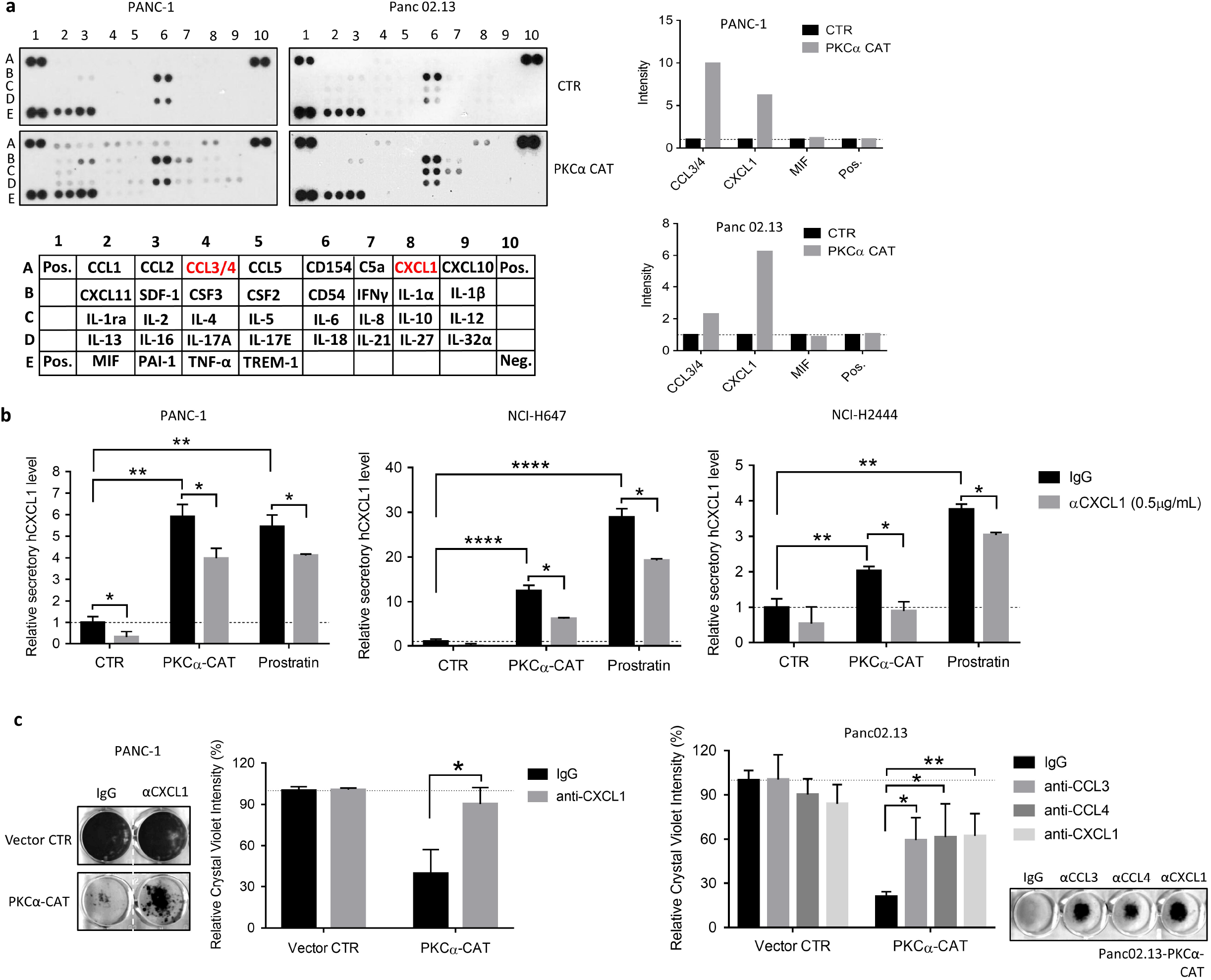
Activation of PKCα induces the expression and secretion of CCL3, CCL4, and/or CXCL1 in KRAS mutant tumor cells. (a) Detection of human cytokines and chemokines by using Proteome Profiler Human Cytokine Array Kit in human pancreatic cancer cells expressing constitutively active form of PKCα. The cells were first transfected with PKCα-CAT, and the total cell lysates were harvested 72 hours post-transfection for the antibody array analysis. The intensity of each dot was determined by ImageJ and normalized by the control dots. Cells expressing empty vector were used as control. (b) ELISA used to detect the secretion of CXCL1 from cells with activated PKCα to the culture supernatant. Cells were treated with prostratin at 2 μM or transfected with constitutively active form of PKCα. Culture supernatant was harvested 72 hours post-treatment or transfection. Cells transfected with empty vector and treated with DMSO were used as control. Prior to ELISA, culture supernatant was normalized with the number of attached cells at the time of harvesting. (n=3; * P < 0.05; ** P < 0.01; **** P < 0.0001) Means are plotted as bars with error bars showing s.d.. (c) Colony formation assay followed by crystal violet staining in the tumor cells expressing active PKCα with treatment of neutralizing antibodies as indicated. Cells seeded in 12 well plates were transfected to ectopically express constitutively active mutant of PKCα, followed by treatment of indicated neutralizing antibody at 0.5 μg/mL till the visible colonies formed. (n=3; * P < 0.05; ** P < 0.01) Means are plotted as bars with error bars showing s.d..

In summary, we discovered that prostratin suppresses oncogenic KRAS-mediated malignancies through the activation of PKCα, which exclusively phosphorylates KRAS at serine 181. Consequently, PKCα acts as a tumor suppressor in the cellular context of mutant KRAS, and, instead of promoting cell death, the activation of PKCα induces senescence through an elevated p21 expression, dependent on PTPN14 which suppresses YAP activity. The activation of PKCα, disrupts KRAS-CaM axis and increases the level of cytosolic CaM thus regulating the nuclear localization of a CAM dependent transcription factor CAMTA1. The KRAS mutant cells undergoing active PKCα-induced senescence exhibit a unique secretory phenotype, specifically an increased production of CCL3/4 and/or CXCL1, whose activity partially mediates the growth inhibition caused by active PKCα. Our results suggest that specific activation of PKCα can be an alternative strategy to target overall oncogenic KRAS-driven human cancers.

## DISCUSSION

Throughout the development of KRAS inhibitors, including the breakthrough G12C inhibitors, most RAS research has focused on genetic or pharmacological disruption of KRAS activity through its G domain, which functionally regulates the molecular GTP/GDP switches, and which contains RAS mutation hotspots. The G domains of all three cancer-associated RAS isoforms are similar, and only the flexible C-terminal HVR can distinguish KRAS from other RAS isoforms. However, whether the HVR domain dictates the oncogenic outputs of KRAS remains incompletely explored. Consistent with our previous report that the binding of KRAS to CaM through serine 181 located in the HVR determines the specific downstream KRAS signaling pathway and CSC-like properties [15], our results here further reveal the specific regulatory mechanism of KRAS phosphorylation resulting in KRAS-CaM dissociation. Our findings significantly contribute to the knowledge of regulating KRAS activity through its HVR domain and can subsequently be used to drive the development of a novel and selective inhibitor of KRAS, which is urgently needed to treat patients suffering from malignancies driven by oncogenic KRAS. Since this inhibitory mechanism functions by targeting the HVR domain of KRAS, such inhibitors while being extremely selective towards KRAS, will not display a narrow range of efficacy against a specific mutant form of KRAS.

The HVR domain of KRAS contains a polybasic region that harbors three potential phosphorylation sites. Among these regions, serine 181 has been shown to be phosphorylated by PKC [17]. Coupled with our previous study, our findings here show that a non-tumor-promoting PKC modulator, prostratin, can specifically block the interaction of KRAS and CaM through activation of PKCα and phosphorylation of KRAS at serine 181. Mark Philips and colleagues proposed a similar concept for the use of PKC agonists, which stimulate phosphorylation of KRAS at serine 181, as a therapeutic approach to treat KRAS-dependent tumors [17]. In their study, treatment with bryostatin I at 1 μM led to phosphorylation of KRAS, and induced apoptosis in NIH 3T3 cells transformed with oncogenic KRAS through rapid dissociation of KRAS from the plasma membrane [17]. On the contrary, we did not observe subcellular translocation of the KRAS protein or apoptosis induced by the treatment with prostratin in tumor cells harboring mutant KRAS. Notably, all PKC-C1 domain agonists exhibit a biphasic dose-dependent effect on activating PKC[67] with maximal activation achieved at 10 nM for Bryostatin I [68, 69]. Thus, the effects induced by a high concentration (1 μM) of bryostatin I in KRAS-dependent tumor cells may not be specifically caused by the activation of PKC isozymes.

Additionally, non-tumor-promoting PKC modulators, including bryostatin I, have shown significant growth-inhibitory activity against various cancer cell lines *in vitro* and *in vivo* [70-73]. Bryostatin family compounds have therefore been examined in phase I and II studies using a wide range of tumor types. However, the results of several phase II studies using bryostatin I alone were disappointing in multiple types of cancers, including colon cancer, of which 30% - 40% carry a KRAS mutation [74, 75]. The unsatisfactory clinical results may be caused by the inconsistent oncogenic or tumor-suppressive roles of PKC isozymes across human cancers. Moreover, we demonstrate here that prostratin but not bryostatin I/II has antitumor activity specifically in cancer cells expressing mutant KRAS, indicating that the clinical applicability of PKC activators as anticancer agents is likely context-specific (i.e., based on the tumor type and driver genetic alterations). Indeed, our results in this study reveal a tumor-suppressive role of PKCα in KRAS mutant cancer cells.

PKC isozymes regulate diverse cellular events, including cell proliferation and protein secretion. In addition, PKC isozymes function in immune cells and further modulate both innate and adaptive immunity. Different PKC isozymes are expressed in and act on various immune cells, such as T cells, B cells, and macrophages [76]. Broad-spectrum PKC activators can therefore cause transient inflammation and rapid release of cytokines (cytokine storm) in the host. Such a drastic immune response can compromise the clinical applicability of PKC activators as anticancer agents. We found that PKCα but not other PKC isozymes specifically phosphorylates KRAS and consequently inhibits KRAS-driven malignancies. Activating or restoring the function of PKCα can be a novel therapeutic avenue for treating cancers driven by oncogenic KRAS with reduced systemic toxicity.

The discovery in the 1980s that PKC is a receptor for the tumor-promoting phorbol esters fueled the dogma that PKC is an oncoprotein (Garg et al., 2014; Kikkawa et al., 1983). Yet, years of clinical trials for cancer using PKC inhibitors not only failed, but in some instances worsened patient outcome. The recent analysis of cancer-associated mutations in the PKC kinase family from diverse cancers revealed that PKC isozymes are generally inactivated in cancer, supporting a tumor suppressive function. [24, 77]. Indeed, prior reports have indicated the potential tumor suppressive role of different PKC isozymes, including PKCα, in some types of cancers. Kazanietz and colleagues reported that PKCα induces senescence in non-small cell lung cancer cells through upregulation of p21 by using two cancer cell lines, H358 and H441 [78]. Alan Fields and colleagues observed that, upon Cre induction, genetic depletion of PKCα worsens the overall survival rate in LSL-KRAS-G12D mice by the activation of the p38 MAPK-TGFβ signaling axis [54]. Their findings have hinted towards a role of PKCα as a tumor suppressor. However, in the study from the Kazanietz group, the upregulation of p21 is a global effect of PMC, which is a potent agonist to activate PKC in an isozyme-unbiased manner. In addition, they suggest that PKCα shows such tumor-suppressive effect during the activation in the S phase, which should be universal to all cancer cells. More specifically, the tumor suppressive mechanism of PKCα from their results does not completely explain whether the tumor-suppressive role of PKCα is specific to KRAS mutant tumor cells. Our study here not only confirms PKCα as a tumor suppressor but also elucidates an alternative mechanism underlying the tumor suppressive effect of activated PKCα specifically in the cellular context of mutant KRAS.

Genetic alterations, including truncating mutation, deep deletion, and loss-of-function mutations, of PKCα rarely occur (< 1%) in patients with PDACs (TCGA, Firehose legacy & PanCancer Atlas); and, therefore, a molecular mechanism that down-regulates maturation and activation is required to cease the signaling activity of PKCα. The dephosphorylation of conventional and novel PKC isozymes by PHLPP1 is a key mechanism to fine-tune the agonist-evoked signaling transduction and the cellular levels of pan-PKC [79, 80]. Upon dephosphorylation, PKC isozymes are subject to ubiquitination and proteasome-dependent degradation [81]. Dephosphorylation of active PKC is therefore a key mechanism to down-regulate PKC. While the regulatory mechanisms underlying the maturation and activation of PKC has been extensively studied [82-84], less information is available on how PKC signaling is terminated both from a global perspective and isozyme-specific differences. Alexandra Newton and colleagues uncovered that the phosphatase PHLPP1 opposes PKC phosphorylation, leading to the degradation of active PKC [80, 85]. Furthermore, in PDACs, they discovered that high PHLPP1 levels lead to low levels of some PKC isozymes including PKCα, which is associated with poor patient survival [85]. Our data on reduced level of phosphorylated PKCα in progressed pancreatic cancer (Extended Data Fig. 5f) partially echoes their findings to indicate that inactivation of PKC could essentially facilitate the tumor progression in cellular context-specific manner (i.e., based on the tumor type and driver genetic alterations such as KRAS mutations).

While somatic mutation of KRAS contributes to numerous types of tumorigeneses, germ-line mutations of KRAS are associated with developmental disorders or RASopathies, including Noonan syndrome [86]. Senescence has been speculated to be a possible contributor to certain complex pathophenotypes of RASopathies [87]. As shown by this study activation of PKCα can be an alternative approach to target KRAS and therefore be a candidate to treat these RASopathies. To facilitate the use of PKCα agonists as an alternative anti-KRAS agent in the clinic, it is necessary to have a biomarker specifically indicating the activation of PKCα and/or the changes in the KRAS/CaM signaling axis. While the activity of PKCα can be determined by phosphorylation of a threonine residue within its activation loop (a-loop), PKCα shares a highly conserved amino acid sequence of a-loop with other two conventional PKC isozymes, PKC-β and -γ. The conserved region across different PKC isozymes increases the cross-reactivity of antibodies and challenges the applicability of using phosphorylated PKCα in the a-loop as a specific biomarker. Our data here suggest that a CaM-dependent transcription activator, CAMTA-1, is stabilized and translocated to cell nucleus in the response to the KRAS-CaM dissociation and, therefore, could be a novel marker to indicate the therapeutic efficacy of PKCα agonists. CAMTAs are highly conserved proteins, are expressed broadly in the nervous system and upon depletion confer pleiotropic behavioral defects in flies, mice, and humans [88]. In addition, CAMTA1 reportedly functions as tumor suppressor to reduce the CSC-like properties in glioblastoma [89]. Even though we found that CAMTA1 is not directly involved in the PKCα-mediated senescence program reported here (data not shown), the nuclear translocation of CAMTA1 induced by PKCα-KRAS-CaM signaling axis might be involved in regulation of some other tumor suppressive function in context of KRAS-driven cancers, such as inhibiting CSC-like properties or rare cases of neurodevelopmental disorders associated with KRAS mutation [90].

In sum, our findings here can provide a clearer understanding of the physiological roles of PKCα in the development of oncogenic-KRAS driven malignancies and lay the groundwork to develop a new class of anti-KRAS treatments, potentially delivering a broad impact to not only treating cancers but also other diseases associated with activating mutations in KRAS.

## MATERIALS AND METHODS

### Cell Lines

All human cancer cell lines were purchased from ATCC and maintained in appropriate culture media at 37°C and 5% CO_2_. PANC-1 cells were maintained in Dulbecco’s Eagles modified medium supplemented with 10% fetal bovine serum (FBS); other cell lines were maintained in ATCC-modified Roswell Park Memorial Institute (RPMI)-1640 medium. The medium was supplemented with 15% fetal bovine serum (FBS) and recombinant human insulin (Gibco 12585-014) for pancreatic cancer cells and with 10% FBS for both human lung and colon cancer cells. NIH 3T3 cells were originally purchased from ATCC. Ras-G12V-transformed NIH 3T3 cells were generated as previously described and maintained in Dulbecco’s modified Eagle’s medium (DMEM) supplemented with 10% FBS. Rasless MEFs[91] were gifts from the NCI RAS Initiative. KP1233 (a murine KRAS-G12D-driven NSCLC cell line) was a gift from Dr. Tyler Jack’s laboratory at the Massachusetts institute of Technology (MIT) and were grown in RPMI-1640 medium supplemented with 10% FBS. The mouse pancreatic cancer cell line iKRAS* [32] was grown in DMEM supplemented with 10% FBS. The human pancreatic epithelial cell line was purchased from Cell Biologics (Cat. No. H6037) and cultured in a complete human epithelial cell medium (Cat. No. H6621).

### Reagents and Antibodies

Prostratin (CAS 60857-08-1) was purchased from Santa Cruz Biotechnology (Cat. No. sc-203422A). Bryostatin I (Cat. No. 2383), bryostatin II (Cat. No. 5237), and PMA (Cat. No. 1201) were obtained from TOCRIS. Trametinib (S2673) and palbociclib (S1116) were purchased from Selleck Chemicals. The following antibodies were used for Western blot analysis or staining in this study: anti-caspase3 (Cell Signaling #9662); anti-cleaved caspase 3 (Cell Signaling #9664); anti-PARP-1 (Cell Signaling #9532); anti-p21 (Cell Signaling #2947); anti-Lamin B1 (Cell Signaling #13435); anti-phospho-CaMKII (Thr286) (Cell Signaling #12716) (Abcam #ab5683); anti-phospho-p44/42 MAPK (Erk1/2) (Cell Signaling #4370); anti-p44/42 MAPK (Erk1/2) (Cell Signaling #4696); anti-KRAS (Sigma #WH0003845M1); anti-β-actin (Sigma #A2228); anti-PKCα (Santa Cruz #sc-8393); anti-CD44 (Santa Cruz #sc-7297); anti-PKCδ (Abcam #ab182126); anti-sox9 (Abcam #ab185966); anti-calmodulin (Abcam #ab45689); anti-HA (Invitrogen #26183 and #PA1-985); anti-pan-RAS (ThermoFisher #MA1-012); anti-phosphoserine (Enzo #ALX-850-023-KI01); anti-CAMTA-1 (ThermoFisher #PA5-103686); anti-CAMTA-2 (ThermoFisher # PA5-103687); anti-PTPN14 (Cell Signaling #13808). The secondary antibodies used for immunofluorescence staining were obtained from Invitrogen (#A32731 and A32723); the secondary antibodies used for Western blot analysis were obtained from LI-COR (IRDye® 800CW Goat anti-Rabbit IgG Secondary Antibody and IRDye® 680RD Goat Anti-Mouse IgG Secondary Antibody) and Sigma (HRP anti-Rabbit or anti-Mouse Secondary antibodies). The constitutively active PKC mutant expression plasmid (PKC alpha CAT) was a gift from Bernard Weinstein (Addgene plasmid # 21234; http://n2t.net/Addgene:21234; RRID: Addgene_21234). YAP/TAZ-responsive synthetic promoter (8xGTIIC-luciferase) was a gift from Stefano Piccolo (Addgene plasmid # 34615; http://n2t.net/addgene:34615 ; RRID:Addgene_34615). pCDNA3-V5-PTPN14-wild type was a gift from Jianmin Zhang (Addgene plasmid # 61003; http://n2t.net/addgene:61003 ; RRID:Addgene_61003). siRNAs (pool) used for knocking down mouse CAMTA-1 were purchased from Horizon (ON-TARGETplus Mouse CAMTA1 siRNA; # L-051054-00-0005), and lipofectamine RNAiMAX reagent (ThermoFisher) was used to transfect siRNA into the target tumor cells. The lentiviral shRNAs against PRKCA and PRKCD were purchased from Horizon (VGH5526-EG5578 and VGH5526-EG5580). Antibodies used for neutralizing the activity of indicated chemokines are: Human CCL3/MIP-1 alpha Antibody (R&D # AF-270-NA); Human CCL4/MIP-1 beta Antibody (R&D # MAB271-100); and Human CXCL1 Antibody (R&D # MAB275-100). Mouse P-PKCα (Serine657) used for IHC is from ThermoFisher (cat. no. PA5-99573).

### Genetically engineered mouse model

Pdx-Cre (B6.FVB-Tg (Pdx1-cre)6Tuv/J; #014647) and LSL-KRAS-G12D (#008179) were originally purchased from The Jackson Laboratory. The number of mice for each breading and genotypes was determined by using Breeding Colony Size Planning Work Sheet, and genotyping was conducted by following the protocol listed in each colony information at The Jackson Laboratory. All the procedures were followed as described in the protocol approved by Univ. Pitt. IACUC (Protocol #: 21089810). The tissues were harvested at indicated endpoint and processed, stained, and graded by the UPMC Hillman Cancer Center Tissue/Histology Core in a double-blind manner.

### Western Blot Analysis

Experimental cells were washed twice in ice-cold phosphate-buffered saline (PBS), lysed in 1% Triton lysis buffer (25 µM Tris [pH 7.5], 150 µM NaCl, 1% Triton X-100, 1 µM EDTA, 1 µM EGTA, 20 µM NaF, 1 µM Na_2_VO_4_, and 1 µM DTT) supplemented with a protease inhibitor cocktail (Roche) and cleared by centrifugation. Protein concentrations were determined by the Bio-Rad Protein Assay (Bio-Rad). Equal amounts of protein extracts were resolved by SDS-PAGE (NuPAGE; Invitrogen), and proteins were transferred to a nitrocellulose membrane. The membrane was immunoblotted with the indicated primary antibodies, incubated with secondary antibodies (details above), and were either visualized with a LI-COR Odyssey scanner or using Immobilon Western Chemiluminescent HRP Substrate (Sigma) and visualized by autoradiography.

### 2D CellTiter-Glo Assay

To measure the short-term effect of PKC activators on cell viability, 2000 cells were seeded in the wells of 96-well plates in a medium containing DMSO, prostratin, or bryostatin I/II as indicated for 72 hours. CellTiter-Glo® 2.0 reagent (Promega) was added and mixed by repeated pipetting. After incubation at room temperature for 20 min, luminescence was measured using a microplate reader. DMSO (the solvent) was used as the control. Relative cell viability was calculated as follows: (luminescence reading for PKC activator-treated cells)/(luminescence reading for DMSO-treated cells) X 100.

### Sphere Formation Assay Quantified by 3D CellTiter-Glo Assay

Cells were harvested, counted, and seeded into Ultra-Low Attachment Culture 96-well plates (Corning Life Science, Cat. No. 3261). One hundred cells were seeded in the wells of low-attachment 96-well plates in medium containing 10% Matrigel® (Corning). The seeded cells and formed spheres were maintained in complete serum containing medium. The medium was supplemented with various PKC activators as indicated until spheres formed in 2-3 weeks. The formed spheres were observed twice weekly and counted one month after seeding.

To evaluate cell viability, an equal volume (100 µl) of CellTiter-Glo® 3D Cell Viability Assay reagent (Promega) was added and mixed by repeated pipetting to improve lysis of spheres.

Plates were incubated for 30 min at room temperature while gently shaking on a rocker. Luminescence was measured using a microplate reader.

### Crystal Violet Staining and Quantification

Cells were seeded in 24 well plates at 1×10^2^ cells per well in complete growth medium with indicated treatment with or without PKC activators. For the effects of constitutively active PKCα in cells, the cells were first seeded at same numbers in 6-well plates. The cells were then transfected with constitutively active PKCα expressing construct at indicated concentrations. 6 hours-post transfections, the cells were cultured back to complete growth medium. After visible colonies formed (usually on the 14^th^ to 21^st^ day depending on the cell line), they were visualized by staining with 0.05% crystal violet (in 0.1% methanol).

For further unbiased quantification of the density of cells adhering to the stained multiwell culture dishes, 1% SDS (300 µl per well of the 24-well plate) was then added to solubilize the stain. The plates were incubated with shaking at room temperature for 30 min or until the color was uniform with no areas of dense coloration in the bottom of the wells. The absorbance of each well was measured at 570 nm.

### CETSA

The CETSA was performed as previously described [42]. In brief, PANC-1 and Panc02.13 cells were treated with the indicated treatments DMSO, prostratin (2 µM), and bryostatin II (0.5 nM) and indicated durations. After treatment, equal numbers of cells were aliquoted in 50 µl of 1X PBS containing a protease inhibitor cocktail in PCR tubes. Using the temperature gradient feature of the thermal cycler, the cell aliquots were treated in parallel at different temperatures as indicated for 3 min. After incubation, another 50 µl of 1X PBS containing protease inhibitors was added, and the cells were lysed by three freeze-thaw cycles (flash freezing followed by thawing at 23°C and vortexing). After lysis, the lysates were centrifuged at 20,000 X g for 20 min at 4°C to remove the insoluble fraction. The soluble fraction was retained, mixed with NuPage™ LDS sample buffer (Thermo Fisher Scientific), and analyzed by Western blotting.

### RBD Pulldown Assay

The RAS pulldown assay was performed using an active Ras pulldown detection kit (Thermo Fisher Scientific) in accordance with the manufacturer’s instructions. In brief, cells were lysed using lysis/binding/wash buffer. Glutathione resin was added to spin cups and precleaned with lysis/binding/wash buffer. The lysates were normalized to obtain equal amounts of total protein between samples. The normalized lysates were incubated with glutathione resin and 80 µg of GST-Raf1-RBD for 1 hour at 4°C with gentle shaking. After incubation, the resin was washed with lysis/binding/wash buffer. Finally, proteins bound to the resin were solubilized in a 2X reducing sample buffer, eluted by centrifugation, and analyzed by Western blotting.

### CaM Pulldown Assay

Cells were washed twice in ice-cold PBS and lysed in 1% TX100-TNM lysis buffer (20 µM Tris [pH 7.5], 5 µM MgCl2, 150 µM NaCl, and 1% Triton X-100) supplemented with 1 µM DTT, 1 µM CaCl2, and protease and phosphatase inhibitors (Sigma-Aldrich). Equal amounts of protein from each sample were added to 50 µl of precleaned CaM binding beads (GE Healthcare, 17-0529-01) in 300 to 500 µl of 1% TX100-TNM lysis buffer and incubated with rotation at 4°C overnight (16-18 hours). The beads were washed 3 times with 1 ml of cold lysis buffer and boiled in NuPage™ LDS sample buffer (ThermoFisher Scientific) to elute the bound proteins. Total cell lysates and pulldown samples were analyzed by Western blotting.

### Senescence-Associated β-Galactosidase Staining

Cells were treated with prostratin, or other compounds as indicated for 7 days to completely induce senescence. To evaluate b-galactosidase activity, cells were stained with a Senescence Cells Histochemical Staining Kit (Sigma, Cat. No. CS0030). The assays were performed following the manufacturer’s instructions. To quantify the population of senescent cells, the positively stained cells from histochemical staining were counted in a double-blind manner (6 images per well were taken and counted), or CellEvent Senescence Green Flow Cytometry Assay Kit was used to detect cellular senescence via β-galactosidase hydrolysis (ThermoFisher, Cat. No. C10840) as per the manufacturer’s instructions. After staining the cells were analyzed on the BD Fortessa flow cytometer to detect green probe fluorogenic signal. Flow data obtained from the cytometer was further analyzed using the FlowJo software.

### Cell Fractionation Assay

Cells were treated with the indicated compounds for indicated duration. After treatment, cells were washed in cold 1X PBS and harvested by scraping in a solution of 1X PBS + 1 mM EDTA. The cell pellets were resuspended in the hypotonic buffer (10 mM Tris-HCI, 1 mM EDTA, and 1 mM DTT supplemented with protease and phosphatase inhibitors). After incubation on ice in hypotonic buffer for 20 min, cells were lysed by repeated passage through a syringe with a 25-gauge 1/2-inch needle. The pellet (nuclear fraction) was isolated by centrifugation at 4°C and 2200 RPM for 3 min. The supernatant was removed and centrifuged at 10,000 RPM for 15 min at 4°C to collect the pellet (crude membrane fraction) and supernatant (crude cytoplasmic fraction). The membrane fraction pellet was washed three times with hypotonic buffer and centrifuged at 10,000 RPM for 10 min at 4°C. The final pellet was resuspended in Western lysis buffer. Membrane contamination from the crude cytoplasmic fraction was removed by centrifugation at 45,000 RPM for 30 min at 4°C using a Sorvall Discovery M120 SE ultracentrifuge with an S120AT2 rotor. The total protein lysates and the membrane and cytoplasmic protein fractions were mixed with NuPage™ LDS sample buffer (Thermo Fisher Scientific) and analyzed by Western blotting. For the nucleus extraction assay, the cells were first transfected with constitutively active PKCα expressing constructs. 48 hours post-transfection, the cellular nucleus was extracted by using Nuclear Extraction Kit (ab113474, Abcam) in accordance with the manufacturer’s instructions.

### PKC Kinase Activity Assay

To measure the working concentration of prostratin or bryostatin required for pan-PKC activation in cancer cells and for use in the functional assays, cells were first treated with compounds as indicated for at least 24 hours. The PKC isoform activities were examined using a PKC enzyme activity kit (Abcam #139437). In brief, cells were washed twice in ice-cold PBS and lysed on ice for 10 min. Then, the lysates were centrifuged at 13,000 RPM for 15 min, and the clear supernatants were analyzed for kinase activity according to the manufacturer’s protocol. Finally, the absorbance of the microplate wells was measured at 450 nm using the Synergy LX Multi-Mode Reader (BioTek).

### Immunoprecipitation

To test whether KRAS acts as a PKC substrate upon stimulation with prostratin or co-transfection of PKCα and oncogenic RAS expressing constructs as indicated, cells were washed 3× with 37°C Hank’s balanced salt solution (HBSS), and the indicated drugs were then added for 30 min in a 37°C non-CO_2_ incubator. Cells were subsequently lysed (1 mM NaF, 1 mM Na_3_VO_4_, 10 nM calyculin A, 1 mM PMSF, 1 mg complete EDTA-free protease inhibitor cocktail [Roche] in RIPA buffer), scraped, collected, and centrifuged at 15,000 X *g* for 30 min at 4°C. The supernatant was collected and incubated with an immunoprecipitating antibody targeting phospho-(Ser) PKC substrates (#2261, Cell Signaling) or HA tag (#71-5500, Invitrogen) for 16–24 hours at 4°C. The lysate-antibody mix was incubated for 3 hours with Protein A/G PLUS-Agarose beads (Santa Cruz) pre-equilibrated in RIPA buffer. After four washes with ice-cold Dulbecco’s phosphate-buffered saline (DPBS), bound protein was eluted by boiling for 10 minutes in NuPage™ LDS sample buffer. Whole-cell lysates, supernatants, and immunoprecipitated samples were subjected to Western blotting with an antibody against KRAS (Clone 3B10-2F2, Sigma, # WH0003845M1), Pan-RAS (#MA1-012, Invitrogen), and HA tag (#26183, Invitrogen).

### Immunofluorescence Staining

Cells were first grown on cover glasses in 24-well plates. After attachment, the cells were treated with PKC activators as indicated for 24 hours. Prior to staining, the cells were washed briefly with 1X PBS, fixed with 4% paraformaldehyde for 15 min, and incubated for 1 min in a solution of 0.01% Triton X-100 in PBS. The cells were then washed three times with 1X PBS and blocked in 1% BSA for 10 min at room temperature. The cells were then incubated with primary antibodies in 1% BSA overnight at 4°C with gentle shaking. Before incubation with secondary antibodies in 1% BSA for 40 min at room temperature in the dark, the cells were washed three times with 1X PBS. The stained cells were counterstained, fixed with ProLong Gold Antifade Mountant with the blue DNA stain DAPI (Thermo Fisher #P36931), and mounted on microscope slides.

### Genome-Wide CRISPR/Cas9 Knockout Screens

In this study, the Human GeCKOv2A CRISPR knockout pooled library was used to identify genes responsible for prostratin resistance in PANC-1 cells. The library was obtained from Addgene (73178-LV). In brief, we first established a stable Cas9-expressing PANC-1 cell line by lentiviral transduction of the Cas9 coding sequence. The transduced cells were selected with 2.0 μg/ml puromycin for 7 days to generate a mutant cell pool, which was then treated with vehicle (DMSO) or prostratin (4 µM) for 21 days. After treatment, at least 3 × 10^7^ cells were collected for genomic DNA (gDNA) extraction to ensure greater than 400X coverage of the GeCKO v2A library. The sgRNA sequences were amplified using NEBNext® High-Fidelity 2X PCR Master Mix and subjected to massive parallel amplicon sequencing carried out by the UPMC Genome Core. Sequencing data from the Cas9 screen were analyzed using MAGeCK (V0.5.9.4, Li et al., 2014) [92] with the “test” option and default settings. Two replicates of untreated and treated conditions were compared, and the “combined gene summary” output, which uses information from the 4 sgRNAs in the library for each gene, was used to compute a per-gene score.

### Establishment of Cell Lines with CRISPR/Cas9-Mediated Gene Knockout

Cancer cell lines with PKCα knockout were established following a previously described protocol [93]. The sequences of the forward/reverse primers for generating PRKCA gDNA were as follows: For1- AGGAAGGAAACATGGAACTC; Rev1- GAGTTCCATGTTTCCTTCCT; For2- CCTTGACCGAGTGAAACTCA; Rev2- TGAGTTTCACTCGGTCAAGG; For3- GCTCCACACTAAATCCGCAG; and Rev3- CTGCGGATTTAGTGTGGAGC. Western blot analysis was used to confirm knockout. At least two different knockout clones were selected for experiments. Cells with Cas9 expression but without knockout were used as control cells.

### Re-expression of PKCα

Cancer cells with PKCα knockout or lacking endogenous expression of PKCα were infected with a lentiviral vector carrying human PRKCA cDNA ORF (Horizon Discovery, # OHS5899-202625016). The infected cells were selected in blasticidin (10 µg/mL) containing complete medium for at least two weeks. The re-stored expression of PKCα in the pool of cells post-selection was confirmed by Western blot analysis.

For the expression of HA-tagged PKCα, the cells were transfected with pHAGE-PKCα-WT, -CAT, or -DN, which were gifts from Dr. Bernard Weinstein (Addgene plasmid # 21232; http://n2t.net/addgene:21232; RRID:Addgene_21232; Addgene plasmid # 21234; http://n2t.net/addgene:21234; RRID:Addgene_21234; Addgene plasmid # 21235; http://n2t.net/addgene:21235; RRID:Addgene_21235, respectively) by using Lipofectamine 3000 (Invitrogen). The ectopic expression of PKCα was confirmed by Western blotting and probing for the HA-tag.

### Tumor xenografts

All experiments were approved by the Institutional Animal Care and Use Committee (IACUC) of the University of Pittsburgh. Human pancreatic cancer cells were subcutaneously injected into nude mice (Nu/Nu) at 0.1 × 10^6^ cells per flank. Palpable tumors were measured once a week. The animals were divided into at least five mice per group. Mice bearing tumors derived from MIA PaCa2 were monitored for one month (two months for mice bearing PANC-1-derived tumors) and euthanized when distressed. All studies were conducted in accordance with the IACUC, and all relevant ethical regulations were followed. Equal numbers of female and male mice were used in the study.

### Detection of active calmodulin in the cytosolic fraction

Cells were first transfected with cytosol-targeted FRET biosensor, which was a gift from Jin Zhang (Addgene plasmid # 64729; http://n2t.net/addgene:64729; RRID:Addgene_64729). The cells were treated with PKC activators 24 hours post-transfection, and the reading of YPet was detected and quantified by using Synergy LX Multi-Mode Reader (BioTak) at different timepoints. To measure the effect of PKCα expression in the cytosolic distribution of CaM, the cells were co-transfected with pcDNA3-Cyto-CaNAR2 and indicated pHAGE-PKCα construct at 1:1 ratio. YPet signal was measured at 48^th^ hour post-transfection, and the numbers of viable cells defined by CellTiter Glo were used for normalization.

### Quantitative PCR

Total RNAs were isolated and purified with the QIAGEN RNAeasy kit; 1 μg RNA per specimen was reverse-transcribed into cDNA with the SuperScript™ First-Strand Synthesis System for RT-PCR (Invitrogen). Possible contamination of genomic DNA was excluded by treatments of DNase I. Quantitative real-time PCR array analysis was performed with SYBR Green (Applied Biosystem). Fold differences and statistical analyses were calculated with the GraphPad Prism 4.00 for Windows (GraphPad Software). The oligo sequences used to detect human CDKN1A are forward 5’ TGTCCGTCAGAACCCATGC 3’ and reverse 5’ AAAGTCGAAGTTCCATCGCTC 3’.

### Luciferase assay

Cells were transfected with YAP/TAZ luciferase reporter construct along with pRL-CMV-Renilla luciferase control reporter vector by Fugene6 (Roche). Luciferase activity was measured with Dual Luciferase Reporter Assay System (Promega; #E1910) according to manufacturer’s instructions. The reporter’s firefly luciferase activity was normalized to the levels of Renilla luciferase used as an internal control reporter. The relative luciferase activity displayed on the Y-axis indicates the ratio between Firefly/Renilla luciferase activities.

### Human Cytokine Antibody Array

The proteome profiler Human cytokine array kit (R&D #ARY005B) was used to detect the level of 36 different cytokines from total cell protein lysates after indicated treatments. The array staining was performed as per the manufacturer’s instructions and resultant autoradiographic films were scanned and analyzed with ImageJ to detect and quantify the intensity of each individual cytokine spot.

### Human CCL3, CCL4, and CXCL1 ELISA

Human CCL3/MIP-1 alpha Quantikine ELISA Kit (R&D # DMA00); Human MIP1 beta ELISA Kit (CCL4) (abcam #ab100597); and Human CXCL1 ELISA Kit (GRO alpha) (abcam #ab190805) were used to detect the indicated chemokines. The assays were performed following the manufacturer’s instructions. To collect the culture supernatant for ELISA, the tumor cells were first treated with 2 μM prostratin or transfected with constitutively active mutant of PKCα. 96 hours post-treatment or transfection, the cell culture supernatant was collected and centrifuged at 3,000 rpm to remove the cell debris. Freshly collected cell-free supernatant was then subjected to ELISA. Each experiment was repeated independently for at least 3 times to get the mean values. The supernatant collected from cells treated with DMSO or transfected with empty vector were used as control.

### Statistical Analysis

Experiments were repeated at least three times when appropriate. Each independent experiment was performed with at least four technical replicates. Data are presented as the mean ± standard error of the mean values. All error bars in these assays indicate the standard deviation (s.d.) of four to eight technical replicates unless noted otherwise. Prism 7.0 software (GraphPad Software Inc.) was used for all statistical analyses, and *P* values < 0.05 were considered statistically significant. Paired or unpaired *t* tests were used for comparisons between two groups where appropriate.

## Supplementary Materials

**Table S1.** Knockdown efficiency and its corresponding effect in senescent population responding to prostratin treatment

| Gene Name | Knockdown efficiency (determined by qPCR) (n=12) |
| --- | --- |
| PTPN14 | 97.11% ± 1.24 |
| EP300 | 91.84% ± 2.1 |
| AHR | 96.1% ± 1.34 |
| MAPK11 | 95.33% ± 1.8 |
| MED12 | 95.33% ± 1.54 |
| LSM12 | 98% ± 1.78 |
| non-targeting |  |

**Table S2.**
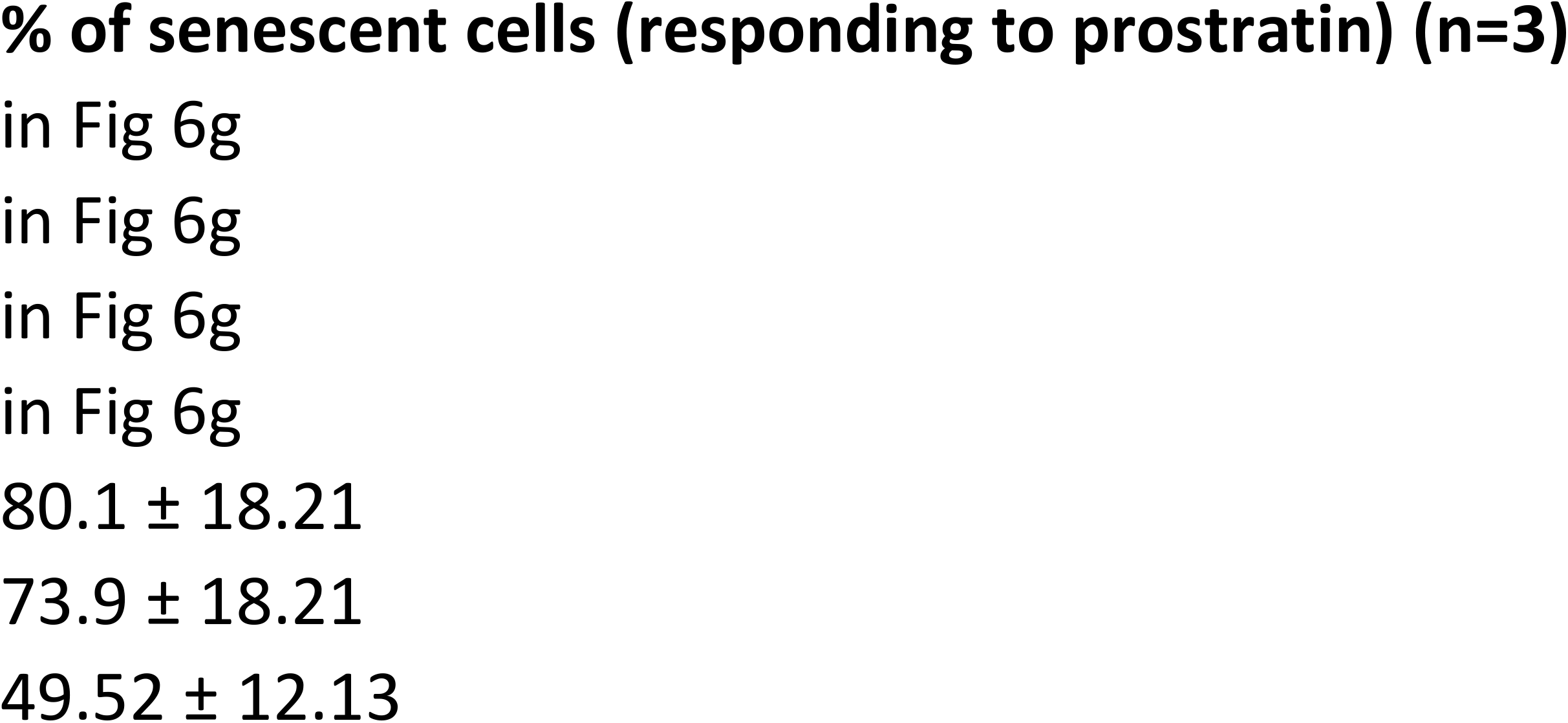
The prediction of CaM binding domains in PTPN14 protein

Data file S2. CRISPR-Cas9 screen gene summary

Data file S1. CRISPR-Cas9 screen gene counts

## Acknowledgments

We thank the generous support from UPMC Hillman Cancer Center Internal Startup funding (M-T Wang).

## Funding

National Institutes of Health grant R21CA259457 (Y.N.G).

## Author contributions

Conceptualization—MTW

Methodology—TJ, YNG, JSO, AU, MTW

Investigation—TJ, AK, ES, MTW

Visualization—TJ, MTW

Supervision—MTW

Writing—original draft: TJ, MTW

Writing—review & editing: TJ, JSO, MTW

## Competing interests

Authors declare that they have no competing interests.

## Data and materials availability

All data are available in the main text or the supplementary materials.

## Supplementary Text

**Supplementary Data Fig 1.**
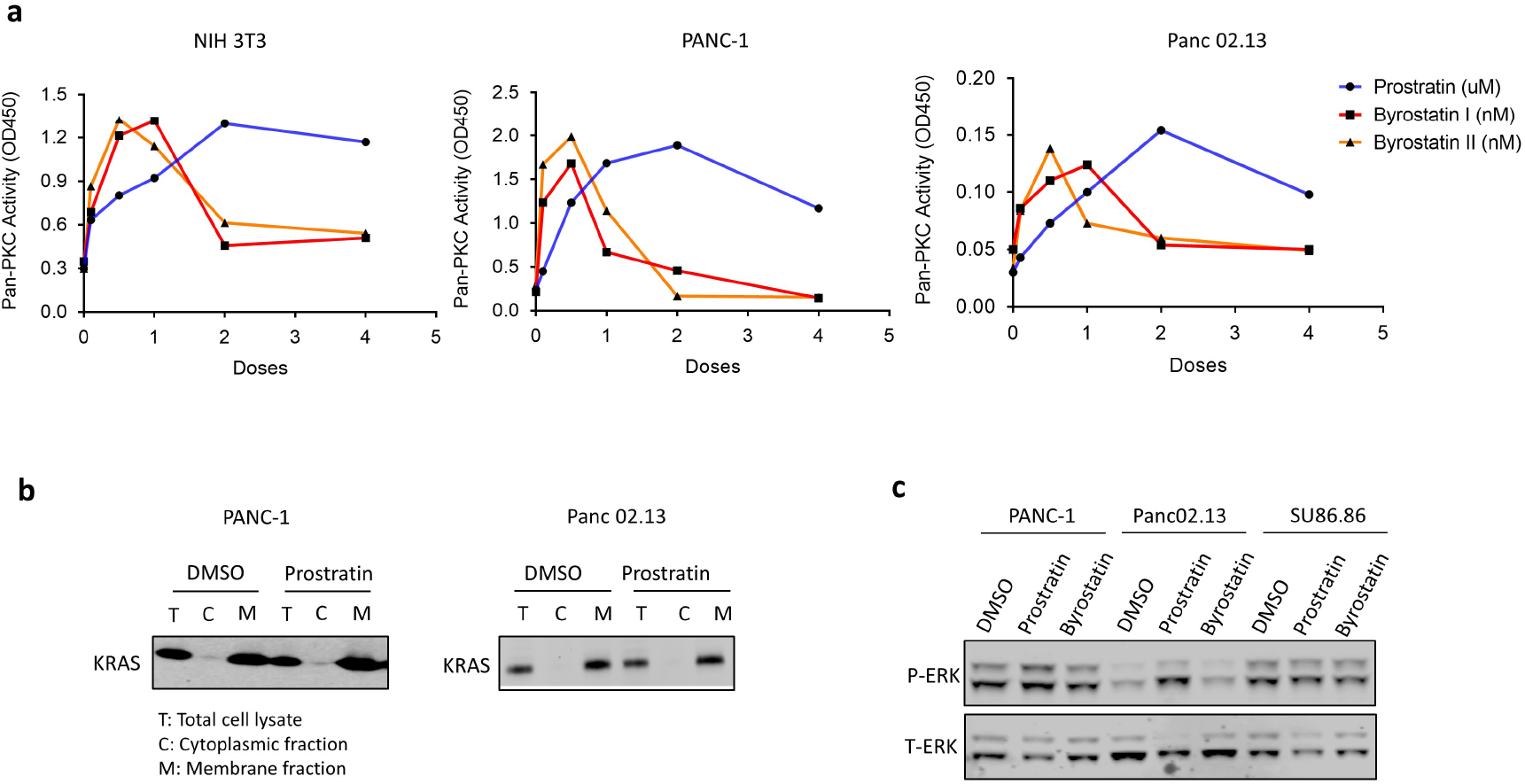
The intracellular working concentration of prostratin to activate PKC, and prostratin does not alter the membrane-localization of KRAS or suppress ERK activation in KRAS mutant cancer cells. (a) PKC enzyme activity assay using pan-PKC substrates to determine the working concentrations of prostratin, bryostatin I, and bryostatin II in NIH 3T3 cells and two different human pancreatic cancer cell lines harboring mutant KRAS. (b) Cell fractionation assay to define the subcellular localization of KRAS in PANC-1 or Panc02.13 cells treated with prostratin at 2 μM for 48 hours. (c) Western blotting for phospho-ERK in cells treated with prostratin at 2 μM or bryostatin II at 0.5 nM for 48 hours. Total ERK was used as the internal control.

**Supplementary Data Fig 2.**
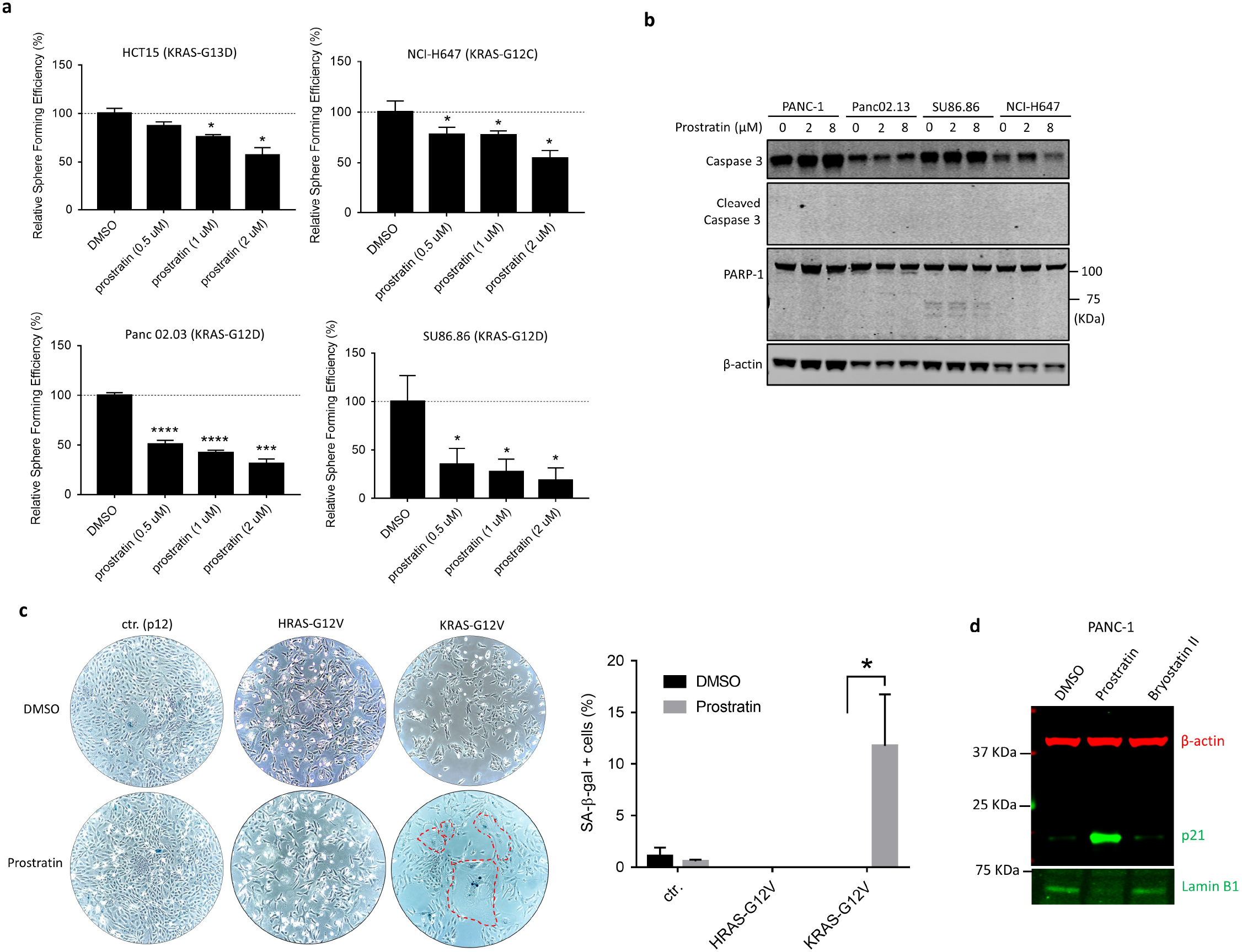
Prostratin suppresses the growth of KRAS mutant tumor cells via induction of cellular senescence. (a) Quantitation of spheres grown in 3D culture in response to treatment with prostratin. The viability of spheres was normalized to that in the control (DMSO-treated) group. (n=6; * P < 0.05, *** P < 0.001) Means are plotted as bars with error bars showing s.d.. (b) Western blot analysis of apoptosis markers in different KRAS mutant cancer cells treated with prostratin for 16-18 hours. (c) β-Galactosidase staining was used to indicate the senescent cells among human pancreatic epithelial cells expressing oncogenic HRAS or KRAS and treated with prostratin at 2 µM for 6 days. The positively stained cells were manually counted in a double-blind manner. Twelve images of each well in a 6-well plate were acquired to calculate the mean value in each group. The error bars indicate the s.d. of three technical replicates. (n=3, *P < 0.05) (d) Western blots for p21, lamin B1, and β-actin in PANC-1 cells treated with prostratin at 2 µM or bryostatin II at 0.5 nM for 7 days.

**Supplementary Data Fig 3.**
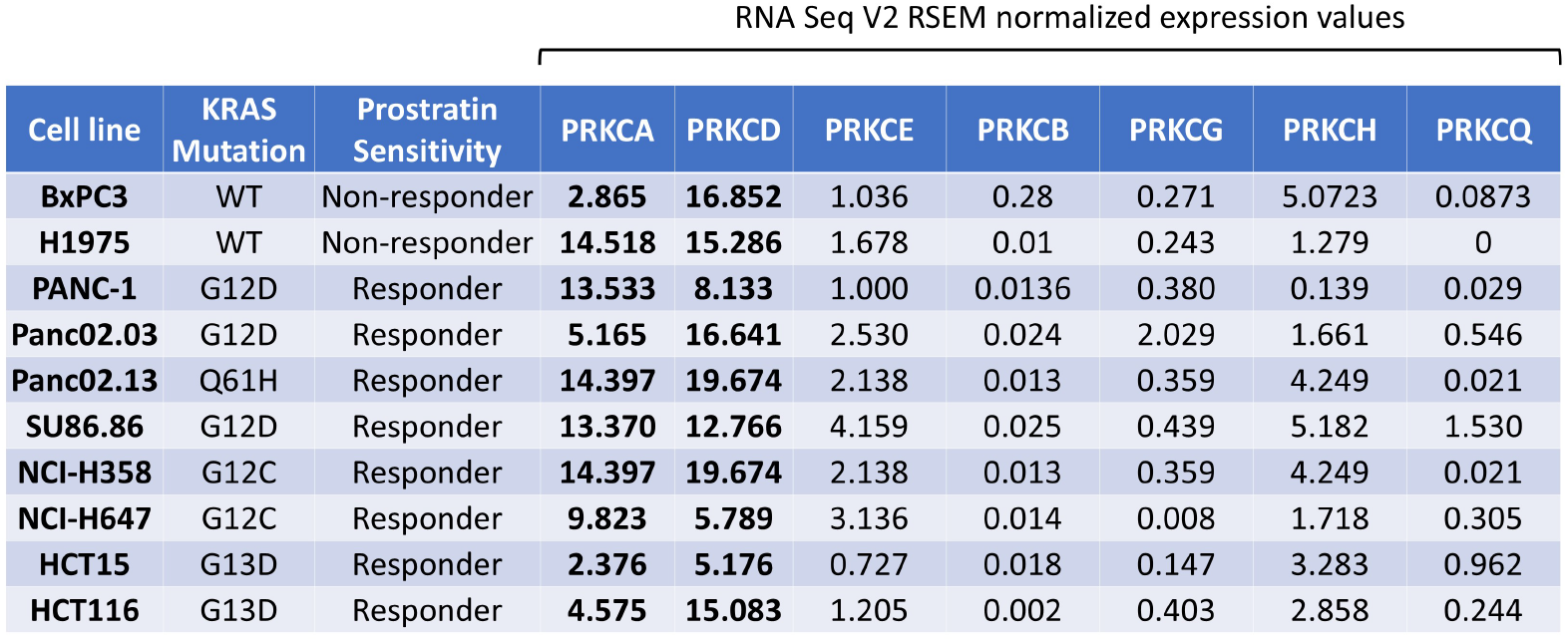
The endogenous PKC expression in different cancer cell lines. Summary of the KRAS mutations, the response to prostratin, and expression level of each PKC isozyme in the cancer cell lines used in this study. mRNA expression levels were obtained from the Cancer Cell Line Encyclopedia (Broad Institute, 2019).

**Supplementary Data Fig 4.**
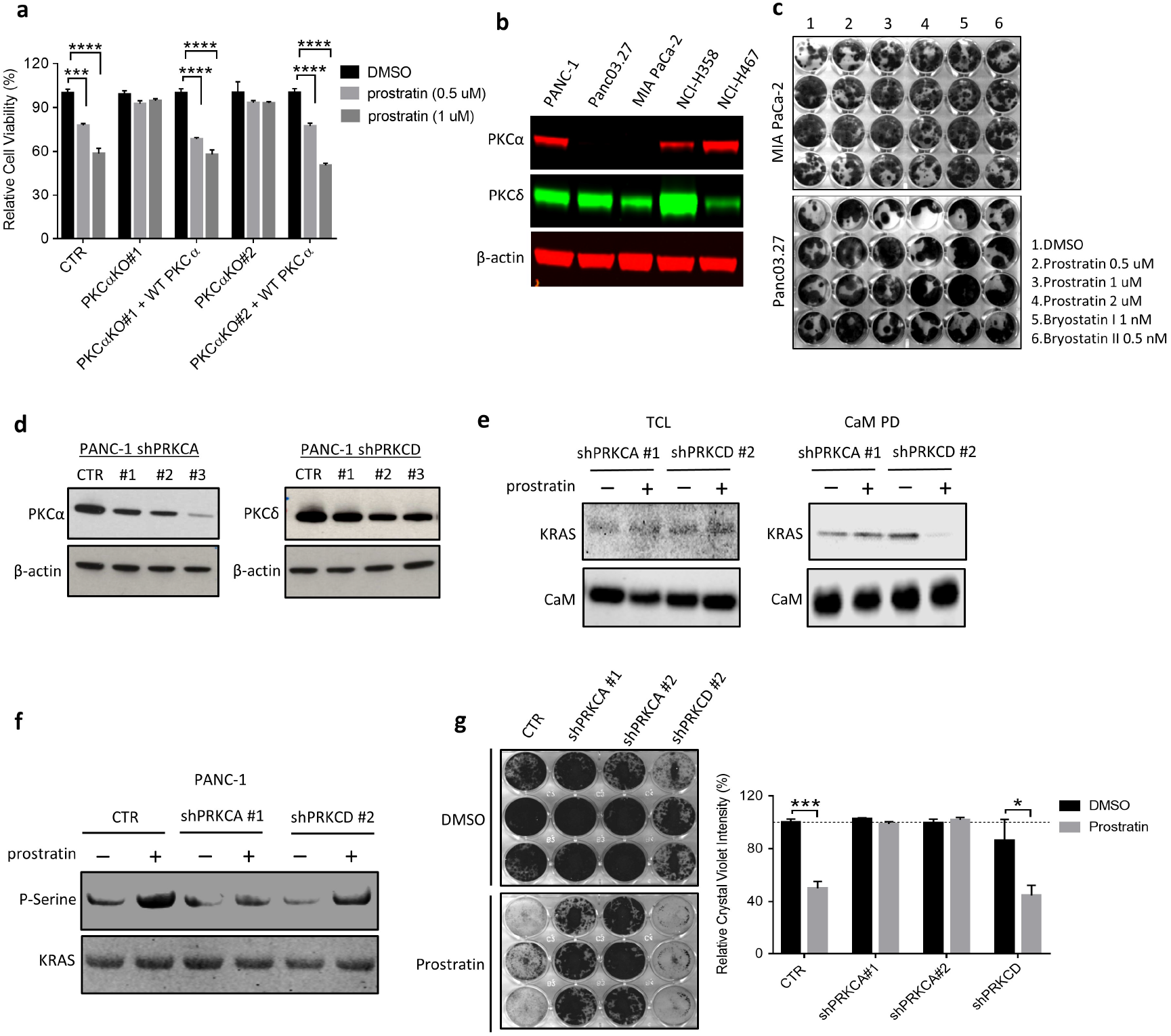
The growth suppression brough by prostratin treatment is only effective to tumor cells harboring mutant KRAS, and PKCδ is not required for the KRAS phosphorylation and growth suppression led by prostratin. (a) CellTiter-Glo assay to determine the viability of PANC-1 cells with modified expression of PKCα after prostratin treatment. Cells were treated for 72 hours. DMSO was used as the control. (n=6, *** P < 0.001, **** P < 0.0001) Means are plotted as bars with error bars showing s.d.. (b) Western blot analysis to evaluate the endogenous expression of PKCα and PKCδ in different KRAS mutant cancer cell lines. (c) Colony formation assay followed by crystal violet staining in MIA PaCa-2 and Panc03.27 cells treated with the indicated compounds. Two hundred cells per well were seeded in complete medium containing the compounds. (d) Western blot analysis confirmed the knockdown of PKCα or PKCδ by different shRNA hairpins in PANC-1 cells. (e) CaM pulldown assay revealed that prostratin treatment caused CaM-KRAS dissociation in PANC-1 cells with knockdown of PKCδ, but not the ones with knockdown of PKCα. The cells were treated with 2µM of prostratin for 24 hours before being harvested for the assay. (f) Ras RBD pulldown followed by Western blotting for phosphoserine in PANC-1. The cells were treated with 2 µM of prostratin for 24 hours before the assay. (g) Colony formation in PANC-1 cells where PKCα or PKCδ has been knocked down by shRNA treated with 2µM of prostratin. Equal numbers of cells (5,000 cells per well) were seeded. (n=3; * P < 0.05; *** P < 0.001)

**Supplementary Data Fig 5.**
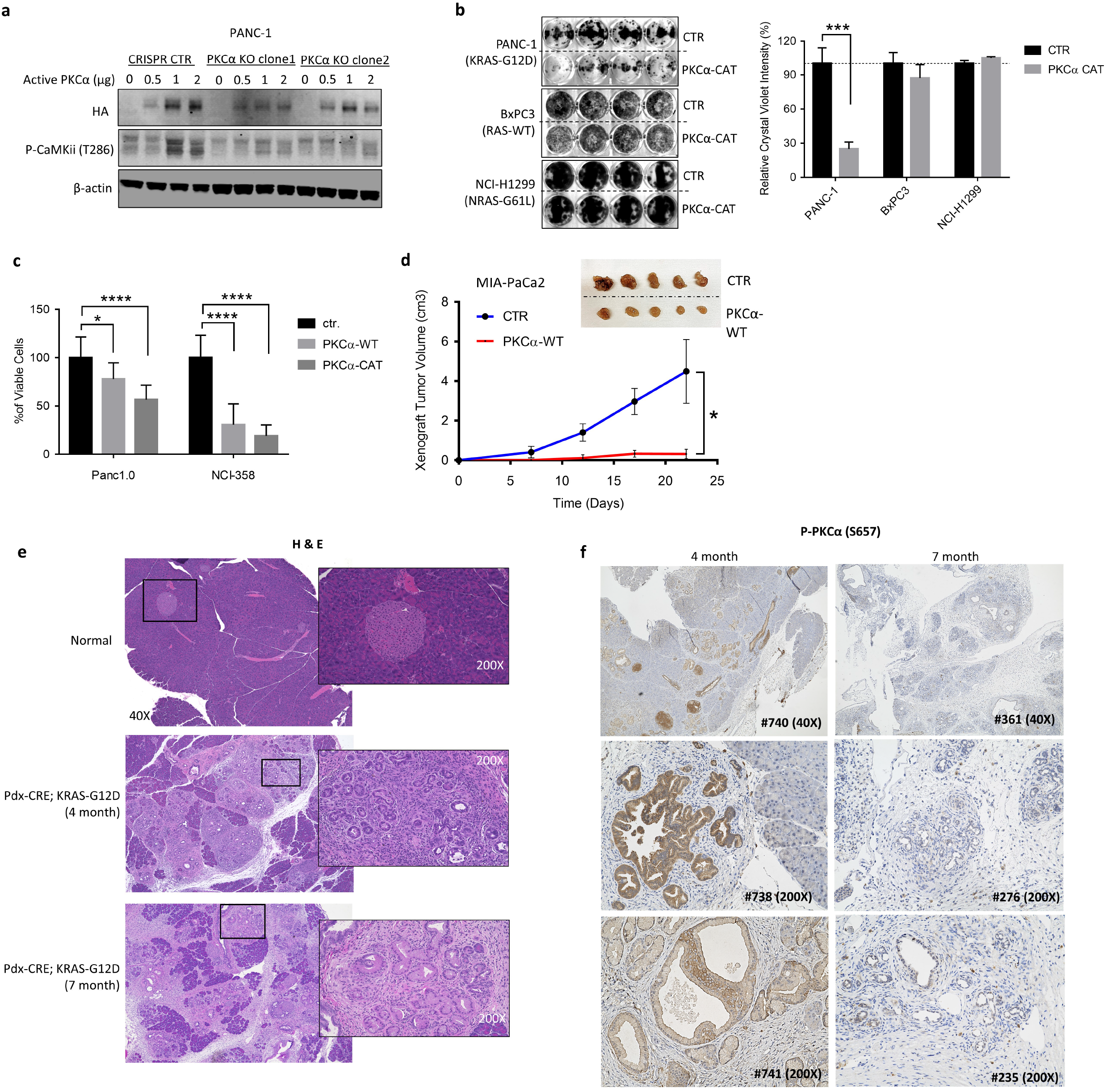
Active PKCα acts a tumor suppressor in oncogenic KRAS-driven cancer. (a) Western blot analysis confirmed the expression level of constitutively active PKCα in PANC-1 cells with knockout of endogenous PKCα. A HA tag was used to indicate specific expression of constitutively active PKCα. Expression of constitutively activated PKCα corresponds with an increase in the phosphorylation level of CaMKII (T286). (b) Colony formation visualized by crystal violet staining in a series of tumor cells with wild type or mutant KRAS which ectopically expressed constitutively active mutant form of PKCα. (n=4, *** P < 0.001) Means are plotted as bars with error bars showing s.d.. (c) Cell viability in the cells expressing wild type of constitutively active PKCα. (n=4; * P < 0.05; **** P < 0.0001) Means are plotted as bars with error bars showing s.d.. (d) Tumor growth curve of xenograft tumors derived from MIA PaCa-2 cells. Error bars here indicate standard error of the mean (SEM). (n=5; ** P < 0.01) (e) H&E staining of pancreatic tumors developed in pdx-Cre; KRAS-G12D mice. The mice were euthanized for pathological analysis at the ages of 4 or 7 months. (f) Pancreatic tumors in pdx-CRE; KRAS-G12D mice stained for phospho-PKCα (S657). The mice were euthanized at the ages of 4 or 7 months (3 mice per group). Each representative image is from an individual mouse. All three mice at 4-month-old had PanIN-1 tumors, and all at 7-month-old had PanIN-3 tumors.

**Supplementary Data Fig 6.**
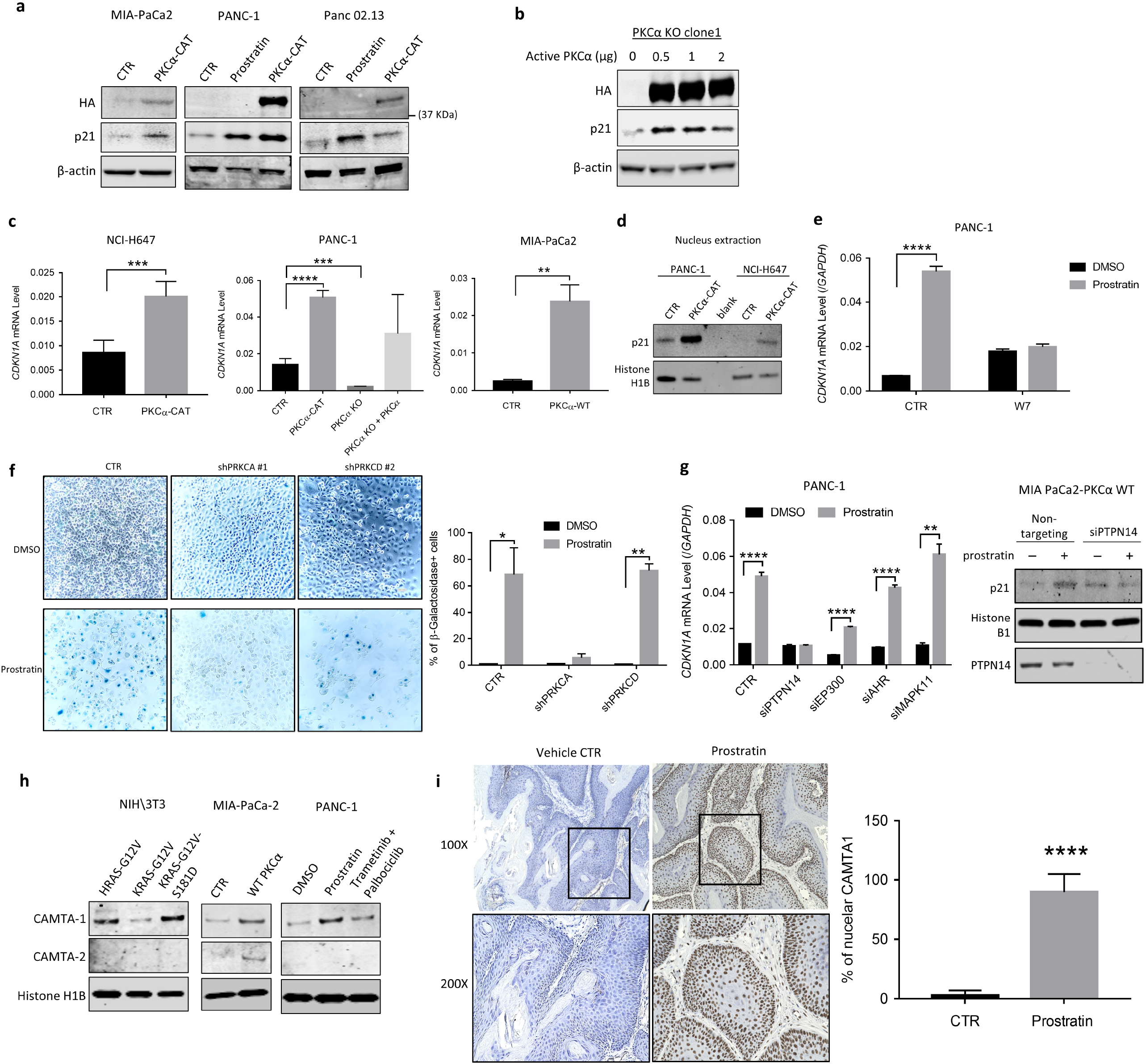
Activated PKCα increases expression of p21 and nuclear CAMTA1. (a) Different pancreatic tumor cells with expression of constitutively active PKCα (HA tagged) or prostratin treatment were subjected to Western blot analysis for p21. HA tag was used to indicate the expression of constitutively active PKCα. (b) Western blot analysis confirmed the expression of HA-tagged PKCα and the increased expression of p21 in PANC-1 cells where the endogenous PKCα had been knocked out by CRISPR/Cas9. (c) Quantitative PCR indicated that the expression level of PKCα affected the expression of p21 (CDKN1A) at mRNA level in tumor cells. (n=3; ** P < 0.01; *** P < 0.001; **** P < 0.0001) Means are plotted as bars with error bars showing s.d.. (d) Nucleus extraction followed by Western blot analysis revealed the increased nucleus-localized p21 level in the cells expressing constitutively active PKCα (PKCα-CAT). Histone H1B was used as the loading control. (e) Quantitative PCR indicated the expression of p21 (CDKN1A) at mRNA level in cells treated with 2 μM prostratin, followed by the treatment of W7 at 25 μM for 72 hours. (n=3; ** P < 0.01; *** P < 0.001; **** P < 0.0001) Means are plotted as bars with error bars showing s.d.. (f) Senescence-associated β-galactosidase staining in PANC-1 cells where PKCα or PKCδ has been knocked down by shRNA. The cells were treated with DMSO (control) or 2µM of prostratin for 5 days before being fixed for staining. (n=4; * P < 0.04; ** P < 0.01) (g) (Left panel) Quantitative PCR indicated the expression level of the expression of p21 (CDKN1A) mRNA in tumor cells where indicated genes were knocked down by siRNA. PANC-1 cells were transfected with siRNA, followed by the treatment of prostratin at 2 μM for 3 days before RNA extraction. (n=3; ** P < 0.01; *** P < 0.001; **** P < 0.0001) Means are plotted as bars with error bars showing s.d.. (Right panel) Nucleus extraction followed by western blot analysis revealed the increased nucleus-localized p21 protein in the MIA PaCa-2 cells expressing wild type PKCα upon the treatment with prostatin at 2μM for 72 hours. Knocking down PTPN14 by siRNA (pool) suppressed the increase in p21 caused by prostratin treatment. Histone H1B was used as a loading control. (h) NIH 3T3 cells transformed by H- or K-RAS-G12V and two different pancreatic tumor cells with overexpressed PKCα or activated PKCα by prostratin treatment were subjected to nucleus extraction followed by Western blot analysis for CAMTA1. Histone H1B was used as a loading control. (i) (Left panel) The representative images of CAMTA1 IHC in mice. Mice bearing KRAS-driven papillomas were treated with vehicle or prostratin, as indicated, for 21 days. (Right panel) Tumor were stained and counted for CAMTA1 in a double-blind manner. Nuclear CAMTA1: control 2.79±4.2%; prostratin 89.7±15.1% (n=4; ***P < 0.001)

**Supplementary Data Fig 7.**
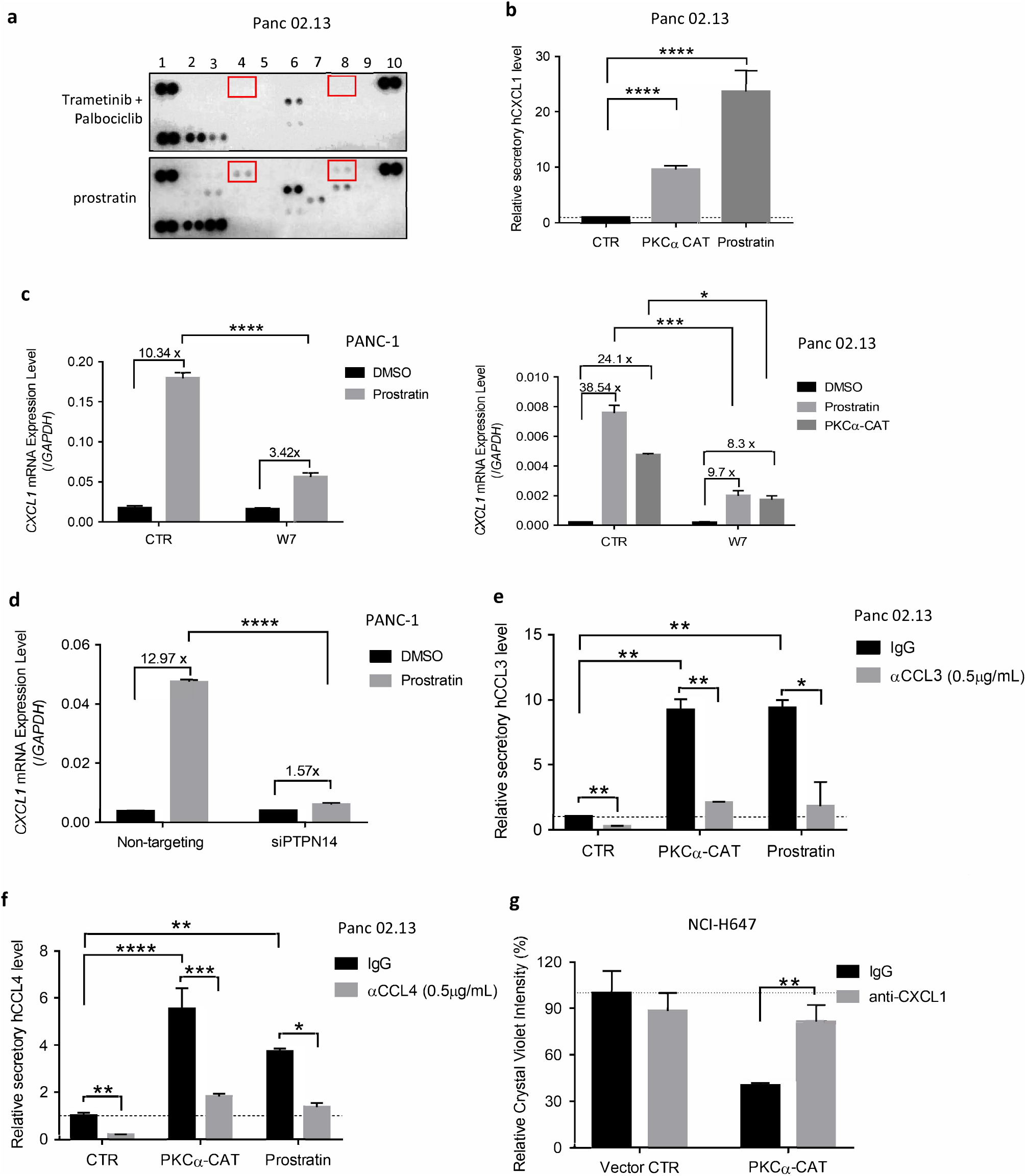
Activation of PKCα induces the secretion of CCL3, CCL4, and/or CXCL1 in KRAS mutant tumor cells in a CaM dependent manner. (a) Detection of human cytokines and chemokines by using Proteome Profiler Human Cytokine Array in Panc02.13 cells treated with prostratin at 2 μM or with a combination of trametinib (10 nM) and palbociclib (0.1 μM) for 72 hours. (b) ELISA to detect the secretion of CXCL1 by Panc02.13 with activated PKCα to the culture supernatant. (n=3; **** P < 0.0001) (c) Quantitative PCR indicated that the expression of CXCL1 at mRNA level in human pancreatic cancer cells treated with prostratin at 2 μM or transfected with PKCα-CAT followed by the treatment of W7 at 25 μM. RNA was extracted 48 hours post-transfection and/or treatment. (n=3; **** P < 0.0001) Means are plotted as bars with error bars showing s.d.. (d) Quantitative PCR indicated that the expression of CXCL1 at mRNA level in PANC-1 cells with siRNA knockdown of PTPN14 and treatment of prostratin at 2 μM. (n=3; **** P < 0.0001) Means are plotted as bars with error bars showing s.d.. (e) – (f) ELISA to detect CCL3 or CCL4 secreted from Panc02.13 with activated PKCα to the supernatant. (n=3; ** P < 0.01; **** P < 0.0001) Means are plotted as bars with error bars showing s.d.. (g) Colony formation visualized by crystal violet staining in NCI-H647 cells expressing active form of PKCα and treated with anti-CXCL1 antibody. (n=4; ** P < 0.01) Means are with s.d. values and plotted as bars with error bars.

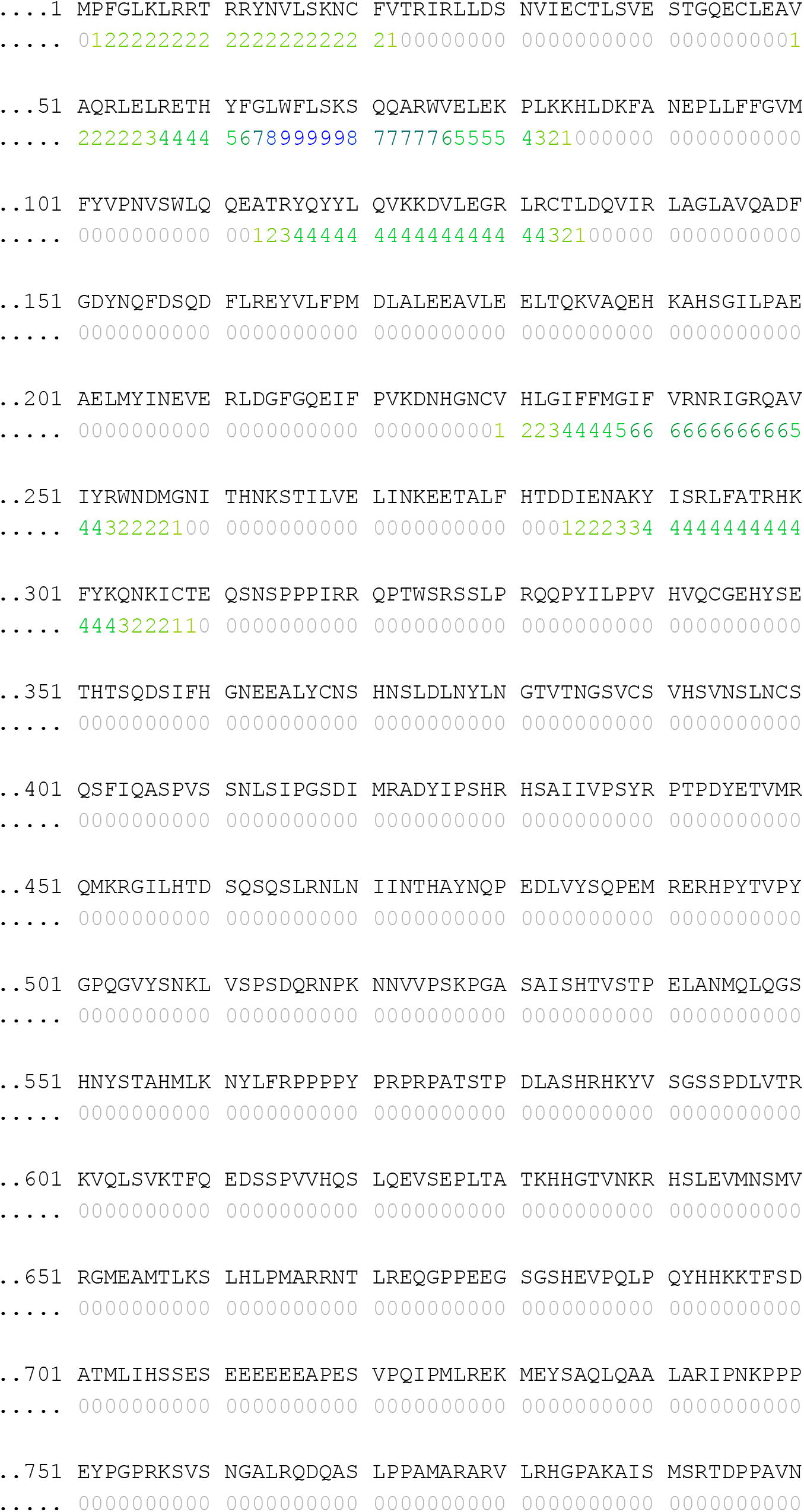

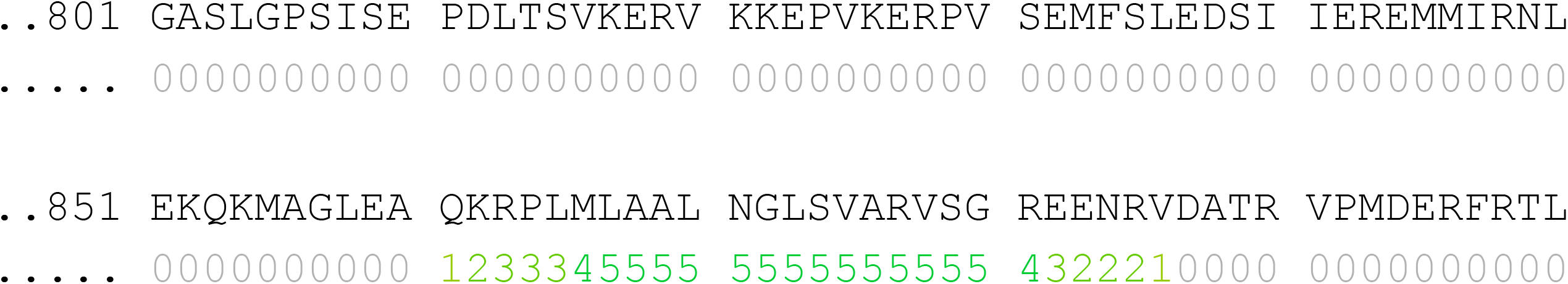

## Notes

### Competing Interest Statement

The authors have declared no competing interest.

### Summary of Updates

1)We have included additional evidence to rule out a role of other cancer-associated PKC isoforms, including PKC-δ and -ε, in the downstream effects of prostratin. 2)We have provided the results to indicate that genetic or pharmacological activation of PKCα induces a cellular senescence program through PTPN14, which suppresses the activity of YAP. 3)We have changed the order of figure 1 and 2 and deleted some data involving the bryostain treatment to provide a better logical flow of the manuscript. These changes do not affect our scientific conclusions.

